# Dissection of acute stimulus-inducible nucleosome remodeling in mammalian cells

**DOI:** 10.1101/573832

**Authors:** Federico Comoglio, Marta Simonatto, Sara Polletti, Xin Liu, Stephen T. Smale, Iros Barozzi, Gioacchino Natoli

## Abstract

Accessibility of the genomic regulatory information is largely controlled by the nucleosome-organizing activity of transcription factors (TFs). Whereas stimulus-induced TFs bind to genomic regions that are maintained accessible by lineage-determining TFs, they also increase accessibility of thousands of *cis*-regulatory elements. Nucleosome remodeling events underlying such changes and their interplay with basal positioning are unknown. Here, we devised a novel quantitative framework discriminating different types of nucleosome remodeling events in micrococcal nuclease ChIP-seq datasets and used it to analyze nucleosome dynamics at stimulus-regulated *cis*-regulatory elements. At enhancers, remodeling preferentially affected poorly positioned nucleosomes while sparing well-positioned nucleosomes flanking the enhancer core, indicating that inducible TFs do not suffice to overrule basal nucleosomal organization maintained by lineage-determining TFs. Remodeling events appeared to be combinatorially driven by multiple TFs, with distinct TFs showing however different remodeling efficiencies. Overall, these data provide a systematic view of the impact of stimulation on nucleosome organization and genome accessibility in mammalian cells.

## INTRODUCTION

Controlled access to the *cis*-regulatory information contained in mammalian genomes is critical for the accurate execution of transcriptional programs. During development, a discrete and largely cell type-specific fraction of a total repertoire of about half a million *cis*-regulatory elements (Dunham et al. 2012) is exposed and thus made available to transcription factors (TFs) and machineries such as RNA polymerases. Such selective exposure of distinct components of the *cis*-regulatory repertoire determines the unique gene regulatory networks in place in different cell types.

Access to the regulatory information in mammalian genomes is controlled by two counteracting forces. On the one hand, many regulatory elements have an intrinsic and DNA sequence-driven propensity to assemble nucleosomes (Tillo et al. 2010; Barozzi et al. 2014), which implies that most functional transcription factor motifs are occluded by default. On the other hand, nucleosome displacement actively enforced by chromatin remodelers enables the selective exposure of the fraction of the regulatory information available for transcriptional control in a given cell type (Barozzi et al. 2014). Such process is instructed by sequence-specific DNA-binding proteins able to recognize motifs in nucleosomal DNA (Zhu et al. 2018) and to instigate nucleosome displacement or remodeling (Zaret and Carroll 2011). Whereas their mode of binding to nucleosomal DNA can be extremely diversified (Zhu et al. 2018), such TFs have been collectively termed “pioneers” and coincide at least in part with lineage-determining TFs, whose expression is triggered by micro-environmental cues in the developmental niche of individual tissues. This is typified by M-CSF (macrophage colony-stimulating factor) induction of the myeloid lineage-determining TF PU.1 in the bone marrow (Mossadegh-Keller et al. 2013). Importantly, the maintenance of nucleosome depletion at most enhancers requires the constitutive binding of lineage-determining TFs, whose depletion results in rapid nucleosome reassembly (Barozzi et al. 2014). A notable exception to this scheme is represented by CpG islands, whose low nucleosome occupancy is imposed by their peculiar nucleotide composition (Ramirez-Carrozzi et al. 2009). Indeed, the relationship between G+C content and nucleosome assembly is typically bimodal, with very low and very high G+C content being both anti-correlated with nucleosome occupancy (Valouev et al. 2011; Fenouil et al. 2012; Barozzi et al. 2014). Overall, the combination of DNA sequence composition and *trans*-acting factors determines both the degree of nucleosome occupancy of each genomic region and the accuracy of nucleosome positioning, and thus eventually the accessibility of the underlying regulatory DNA (Lai and Pugh 2017).

Importantly, while at some genomic sites nucleosomes are reproducibly maintained at a fixed position, at others they tend to occupy a range of overlapping yet distinct positions in different cells of the same population (Lai et al. 2018), thus resulting in fuzzy signals in micrococcal nuclease (MNase)-based population analyses. In general, compared to nucleosomes at inert genomic regions, nucleosomes at active *cis*-regulatory elements tend to be more reproducibly positioned across different cells in the same population (Lai et al. 2018). Maintenance of well-positioned nucleosomes involves a combination of DNA sequence determinants favouring or disfavouring nucleosome assembly and trans-acting factors, namely TFs working in concert with chromatin remodelers to restrain lateral nucleosome movements (Tillo et al. 2010; Valouev et al. 2011) (Barozzi et al. 2014). The eviction or the shifting of well-positioned nucleosomes therefore requires that such *cis*- and *trans*-acting determinants be over-ruled, while less constrained nucleosomes might provide opportunities for more rapid and less energy-demanding remodeling. Finally, from a biological point of view, inaccurate positioning may generate opportunities for stochastic exposure of the underlying DNA sequence and thus for TF binding and gene activation. Techniques such as ATAC-seq (Buenrostro et al. 2013b) and DNaseI-seq (Boyle et al. 2008) enable the exploration of the accessible, nucleosome-depleted genome and are thus informative of how the regulatory information is exposed in different cells or conditions. However, these techniques do not provide direct information on the nucleosomal changes associated with (and in fact determining) variations in the accessibility of the genomic DNA in different conditions and time windows, from developmental transitions to acute responses to micro-environmental stimuli.

When considering acute responses to stimulation, a general paradigm inferred from the analysis of a wealth of genomic data, is that stimulus-regulated TFs (such as NF-kB and estrogen or androgen receptors) bind to genomic regions that are constitutively occupied and kept accessible by lineage-determining TFs such as PU.1 in myeloid cells and FOXA1/2 in cells of endodermal origin (Ghisletti et al. 2010; Heinz et al. 2010; Wang et al. 2011; Barozzi et al. 2014; Glass and Natoli 2016; Monticelli and Natoli 2017). However, this simple scheme does not account for the common observation that acute stimulation increases accessibility of hundreds to thousands of *cis*-regulatory regions (Mueller et al. 2017; Park et al. 2017), which raises several important questions: *i*) how are nucleosomes associated with *cis*-regulatory elements affected by an acute stimulus? *ii*) How is the pre-existing organization restored after stimulus termination? *iii*) Is the pre-existing organization of nucleosomes at rapidly inducible enhancers designed to enable acute changes in accessibility? *iv*) Are individual TFs endowed with a different ability to disrupt nucleosome organization both qualitatively (e.g. at distinct classes of *cis*-regulatory elements) and quantitatively (with different efficiency)?

To address these questions, in this study we investigated the interplay between DNA accessibility and nucleosome organization on a genome-wide scale in both basal and stimulated conditions by combining ATAC-seq and ChIP-seq on nucleosomal, MNase-digested chromatin from basal and LPS-treated macrophages. We devised and implemented a quantitative framework to detect and classify remodeling events and applied it to determine which changes in nucleosome organization occur at regions showing alterations in either accessibility or TF binding upon stimulation. By integrating these data with a comprehensive panel of time-resolved ChIP-seq profiles for several TFs, we inferred the nucleosome remodeling potential of various TFs regulated by stimulation.

## RESULTS

### The nucleosomal landscape of unstimulated macrophages

Nucleosome preparations were obtained from formaldehyde-fixed untreated or LPS-treated (30’, 60’, 120’ and 240’) mouse bone marrow-derived macrophages. MNase digestion was calibrated to generate a prevalence of mono-nucleosomes over di-nucleosomes (*ca.* 80% and 20%, respectively) (Barozzi et al. 2014), with negligible amounts of poly-nucleosomes. Two biological replicates were used as source of chromatin for ChIP assays with antibodies recognizing histone modifications preferentially associated with poised and active promoters (H3K4me3), poised putative enhancers (H3K4me1) and active *cis*-regulatory elements (H3K27Ac) (**Suppl. Table 1**). MNase-ChIP-seq samples were sequenced to an average depth of 42M reads. A high-depth (800M reads) MNase-seq data set previously generated in untreated macrophages (Barozzi et al. 2014) was used as reference for comparisons. To estimate the nucleosome midpoints and for data visualization, the minus strand coverage was subtracted from the plus strand coverage as described (Haberle et al. 2014). For each nucleosome, the midpoint coincides with the transition between the plus and the minus strand signals.

Compared to MNase-seq, isolation of covalently modified nucleosomes by MNase-ChIP-seq resulted in a consistent enrichment of reads coverage at both promoters and intergenic enhancers (log2-enrichment range: 1.8-4.9; **Figure 1A-B**). Patterns of nucleosome positioning in the two MNase-ChIP-seq replicates were highly reproducible, **Suppl. Fig. 1A-B**).

**Figure 1.**
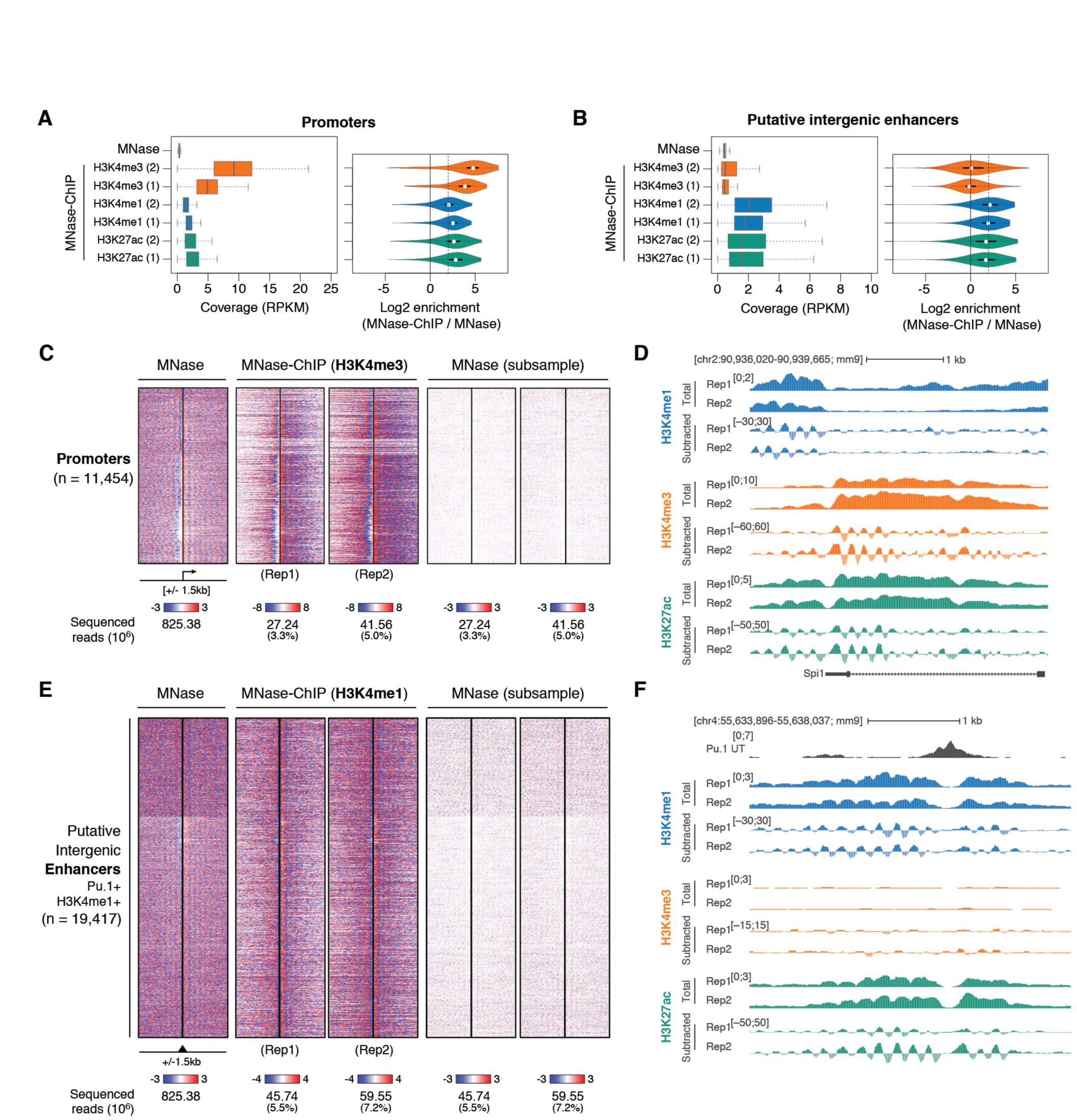
Analysis of nucleosome positioning and occupancy by MNase-ChIP-seq. A-B) Normalized coverage of H3K4me3, H3K4me1 and H3K27ac MNase-ChIP-seq data at promoters (A) and putative intergenic enhancers (B). Violin plots (right) show the MNase-ChIP-seq signal enrichment relative to MNase-seq (log2; dashed line: 4-fold enrichment). RPKM = reads per kb per million mapped reads. C) Nucleosome organization at active promoters. Genomic regions were centered on the main mapped TSS, extended by 1.5 kb on each side and clustered based on H3K4me3 MNase-ChIP-seq signal in the central region (± 150 bps). For comparison, heatmaps based on total and downsampled MNase-seq data are shown. D) Total and subtracted signals from a representative TSS-proximal region. E) Same as (C) but centered on putative extragenic enhancers, using H3K4me1 MNase-ChIP-seq data and centering on the summit of the PU.1 peak, extended by 1.5 kb on each side. F) Total and subtracted signals from a representative TSS-distal, putative enhancer region.

Nucleosome maps at 11,454 H3K4me3-positive promoters and 19,417 H3K4me1- and PU.1- positive putative intergenic enhancers are shown in **Figures 1C** and **1E.** Representative snapshots are illustrated in **Figures 1D** and **1F** and an extended genomic region is shown in **Suppl. Fig. 1F**. Since PU.1 plays a central role in the organization of macrophage-specific enhancers where it is also required to maintain nucleosome depletion (Barozzi et al. 2014), the summit of PU.1 peaks was used as a proxy for the identification of enhancer core regions. Therefore, enhancer maps were centered on PU.1 peak summit (± 1.5 kb) while promoter maps were centered on the main annotated transcription start site (TSS ± 1.5 kb).

Together, these results indicate that the relatively shallow sequencing of MNase-ChIP-seq libraries yielded high-resolution and reproducible nucleosome positioning maps at *cis*-regulatory elements in a mammalian genome (**Figure 1C-F** and **Suppl. Table 1**).

### Two classes of enhancers with distinct nucleosomal symmetry

Visual exploration of the data hinted at the existence of two distinct types of enhancers distinguished by their nucleosomal symmetry, namely by the relative enrichment of H3K4me1-positive nucleosomes upstream and downstream of the enhancer core. Since enhancers are thought to act independently of their orientation, the existence of asymmetric enhancers was intriguing and warranted additional investigations. To systematically classify these two types of enhancers, we computed the ratio between H3K4me1 signals upstream and downstream of the enhancer core regions using different window sizes (**Figure 2A**) and classified as asymmetric those enhancers with an absolute ratio ≥ 1.5 in both replicates. The correlation between upstream and downstream signals among replicates was consistently high (SCC = 0.68-0.74) and was further confirmed in MNase-seq replicates (SCC = 0.77; **Suppl. Fig. 1C-E**). Although the exact number of asymmetric enhancers depended on the window size used for signal summarization, 19-35% of enhancers were consistently classified as asymmetric by our analysis using either H3K4me1 or H3K27ac (**Figure 2A-B**). By comparison, 49-62% of promoters were found to be asymmetric based on H3K4me3 signals (**Figure 2A**). These results were not affected by differences in repeat content or sequence mappability (**Suppl. Fig. 2A-C**).

**Figure 2.**
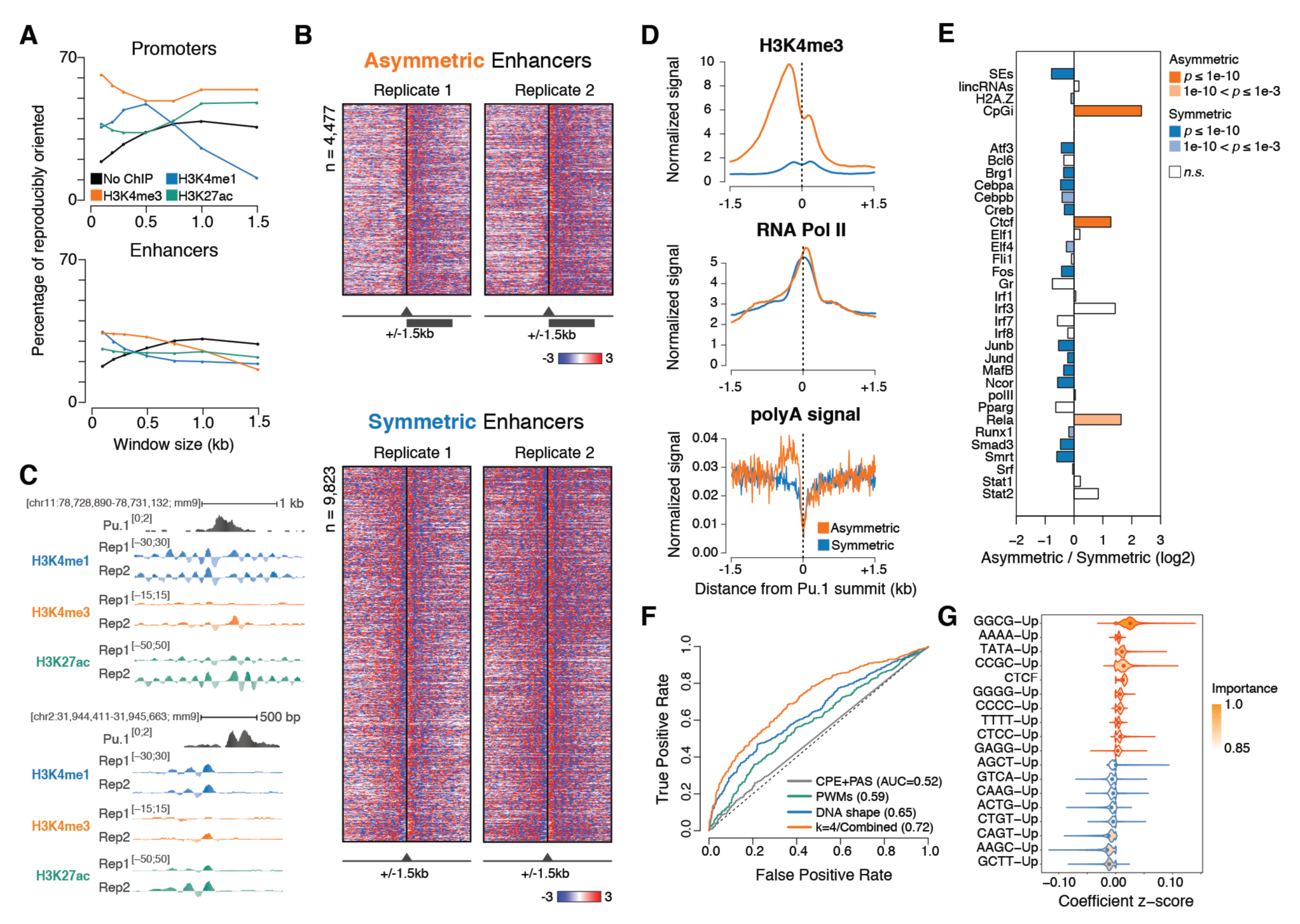
Identification of two classes of enhancers with distinct nucleosomal symmetry. A) Percentage of reproducibly oriented elements, defined as showing concordant normalized signal bias with respect to the core element in both replicates, for both MNase-seq and MNase-ChIP-seq targeting the indicated modified nucleosomes. B) Asymmetric (top) and symmetric (bottom) H3K4me1 MNase-ChIP-seq signals at enhancers centered on the summit of PU.1 peaks. Genomic regions were clustered based on signals in the central region (± 500 bps). C) Representative genomic regions showing a symmetric (top) and an asymmetric (bottom) enhancer. D) Cumulative distributions of H3K4me3, RNA Pol II and polyA signals at symmetric and asymmetric enhancers. E) Enrichment of selected genomic features at symmetric and asymmetric enhancers. *p*-values are from a two-sided Fisher’s Exact Test. F) Receiver operating characteristic (ROC) curve and mean area under the ROC curve (AUC) values for lasso logistic regression models trained on the indicated sets of features. The ROC curve closest to the mean AUC from 100 models is shown. G) Top-ranked features selected by bootstrap-lasso for the combined model in (F). Violin plots are color-coded according to coefficient signs.

Overall, when using a window size of ± 500 bp relative to the enhancer core, 4,477 enhancers (32.7%) exhibited an asymmetric distribution of H3K4me1 nucleosomes while 9,823 (67.3%) were symmetric (**Figure 2A-B** and **Suppl. Table 2**). Representative genomic regions are shown in **Figure 2C**. Compared to symmetric enhancers, asymmetric enhancers shared higher similarity with promoters, including higher levels of H3K4me3, moderate enrichment for CpG islands and CTCF sites (*p* < 1e-10; Fisher’s Exact Test), and an enrichment for poly-adenylation sites (PAS) opposite to the side exhibiting well-organized nucleosomes and RNA Pol II occupancy (**Figures 2D-E** and **Suppl. Fig. 2D**). The two groups of enhancers showed no or minimal differences in terms of evolutionary conservation, relation to nearest active genes and distance from TAD (topologically associating domains) boundaries.

These observations prompted us to test whether DNA sequence features could discriminate symmetric from asymmetric nucleosomal patterns at enhancers. To this end, we considered three sets of features: *i*) *k*-mers frequencies (with 2 ≤ *k* ≤ 4); *ii*) DNA shape features (Chiu et al. 2016) and *iii*) motif scores from a curated collection of >1700 TF motifs (Diaferia et al. 2016). These features were used alone or in combination to train lasso logistic regression classifiers (Comoglio et al. 2015; Comoglio et al. 2018) and prediction accuracies were evaluated on an independent test set. This analysis revealed that TF motifs were only marginally predictive for nucleosomal symmetry at enhancers (mean area under the receiver operating characteristic curve (AUC) = 0.59) and that DNA shape features (AUC = 0.65) and particularly 4-mer sequences (AUC = 0.72) were able to achieve more accurate predictions (**Figure 2F**). In addition, models combining these feature sets did not outperform models solely based on 4-mers (AUC = 0.72), indicating a high feature redundancy. To identify the most predictive sequence features, we performed feature selection on the combined model using bootstrap-lasso, a procedure that assigns high importance to indispensable features (Comoglio and Paro 2014). We found that GC-rich, polyA and TATA sequences upstream of the enhancer core, along with CTCF motifs at the site, were most predictive for asymmetric enhancers, whereas GCTT, AAGC and CAGT sequences were predictive for symmetric nucleosomal patterns (**Figure 2G**).

Together, these results indicate the existence of two distinct classes of enhancers distinguished by the symmetry of their nucleosomal patterns. Moreover, they suggest that such distinctive patterns are primarily determined by DNA sequence features. Since asymmetric enhancers exhibit promoter-like features and enhancers are a major source of long non-coding RNAs (Hon et al. 2017) in addition to bidirectional short RNAs (Kim et al. 2010), these data hint at the possibility that asymmetric *vs*. symmetric nucleosomal organization at enhancers might correlate with their ability to generate different types of transcripts.

### A quantitative framework to measure dynamic nucleosomal changes

To analyze LPS-induced changes in nucleosomal organization at promoters and enhancers, we first devised a quantitative approach aimed at detecting different types of remodeling events. This approach exploits *quantitative changes* in MNase-ChIP-seq signals associated with individual nucleosomes as well as *local changes in correlation* between coverage profiles across conditions (untreated and multiple time points of LPS stimulation) to detect: *i*) shifts in nucleosome position; *ii*) nucleosome evictions and *iii*) locally increased nucleosome occupancy (**Figure 3A**). In the interpretation of these data it should be noted that many inducible genes (such as *IL12b* and interferon-stimulated genes, ISGs) are activated only in a fraction of the LPS-stimulated cells (Weinmann et al. 2001; Shalek et al. 2013). Because of this heterogeneity, remodeling events occurring in a small fraction of cells will be diluted by unaffected nucleosome, thus reducing sensitivity. Therefore, the events identified by our analysis likely underestimate the set of remodeling events occurring at LPS-induced genes.

**Figure 3.**
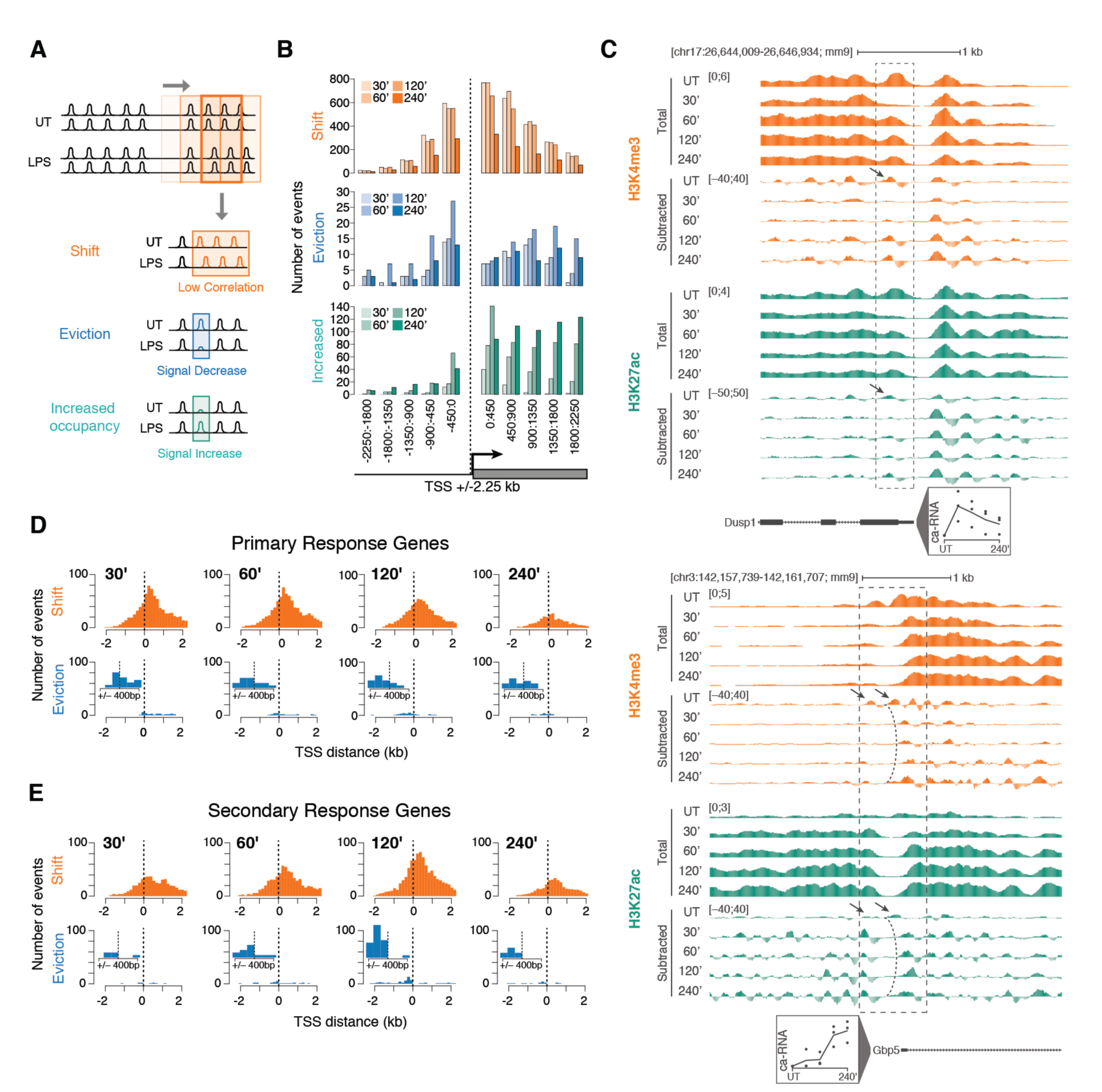
A quantitative approach for detecting inducible nucleosome remodeling events. A) Schematic representation of the approach used to detect different types of nucleosomal changes induced by stimulation. A sliding window of 450 nucleotides (step size 150 bp) was used. Nucleosome shifts were identified as significant drops in correlation between profiles, while evictions and locally increased occupancy were identified as significant decrease/increase that were lost when considering a 10x larger window (4.5 kb). B) Distribution of the three distinct types of nucleosome remodeling events detected at the TSS of LPS-inducible genes at different time points after LPS stimulation. Events are shown in ten consecutive windows of 450 bp each. C) Representative genomic regions showing remodeling of TSS-proximal nucleosomes. D-E) Distributions of nucleosome remodeling events centered on the TSS of primary (D) and secondary (E) response genes at different time points after LPS stimulation. Insets show an enlargement of the distributions of eviction events throughout the central 900 bps around the TSS.

### Nucleosome remodeling at stimulus-regulated promoters

We first applied our analytical framework to the promoter regions of LPS-inducible genes to identify nucleosome remodeling events otherwise undetectable by a conventional ChIP-seq analysis. We used a sliding window of 450 bp to analyze 4.5 kb regions centered on the TSS and detected remodeling events at each time point of LPS stimulation (**Figure 3B**). Remodeling events identified downstream of the TSS often extended further downstream into the body of LPS-inducible genes, likely reflecting the remodeling activity of the elongating RNA Pol II. Conversely, remodeling events upstream of the TSS were generally limited to a proximal region spanning a few nucleosomes. Nucleosome shifts were by far the most common events, while evictions were uncommon. As an example, the *Dusp1* gene is constitutively marked by H3K4me3 and is rapidly induced by LPS, with a peak of transcription at 30’ after stimulation, as indicated by nascent transcript analysis (**Figure 3C**). The overall H3K4me3 signal intensity was only moderately affected by stimulation. However, the +1 nucleosome underwent a near-complete eviction within 30’ of LPS stimulation, followed by a progressive gain in signal that correlated with the gradual reduction in transcriptional activity (**Figure 3C**). At the *Gbp5* gene promoter, one H3K4me3-positive nucleosome upstream of the TSS (left arrow) was evicted, while the downstream nucleosome underwent a sustained shift (**Figure 3C**, bottom panel).

The frequency of remodeling events peaked between 30’ and 60’ at primary response genes (PRGs, **Figure 3D**), while it peaked at 120’ post-stimulation at secondary response genes (SRGs, **Figure 3E**). This result is consistent with the average timing of induction of the genes in the two groups (Bhatt et al. 2012). Similar profiles were obtained by dividing inducible genes into two kinetic classes (early *vs*. late induction) (**Suppl. Fig. 3A-B**). While drops in correlation were detected on both sides of the TSS (with a higher frequency downstream of the TSS), evictions were by far more common upstream of the TSS of secondary response genes (**Figure 3E**).

Overall, when considering a conservative window of 900 bps centered on the TSS, we detected H3K4me3-positive nucleosome remodeling events at the promoters of 437 out of 1,879 LPS-inducible genes (**Suppl. Table 3**). Previous studies demonstrated that LPS-inducible genes with a CpG island in their promoter, which tend to be activated earlier in the LPS response, do not require chromatin remodeling for activation due to the poor propensity of very high G+C content regions to assemble stable nucleosomes (Ramirez-Carrozzi et al. 2009). Conversely, the induction of genes with lower G+C content, which tend to show slower activation kinetics, is dependent on Swi/Snf-mediated nucleosome remodeling (Ramirez-Carrozzi et al. 2009). Therefore, we divided LPS-induced genes from seven increasingly slower kinetic classes (Bhatt et al. 2012) based on the presence or absence of a CpG island and analyzed chromatin remodeling events within each class (**Figure 4A-B**). Nucleosome evictions, which preferentially occurred upstream of the TSS (**Figure 3D-E**), were nearly exclusively observed at the promoters of slowly inducible non-CpG island genes (classes 5-7), with the exception of class 4 (**Figure 4A-B**). Conversely, nucleosome shifts were detected across all classes and likely reflected remodeling events associated with RNA Pol II recruitment and elongation (note that low G+C promoters in class 1, 2 and 3 – the only groups showing limited enrichment – include only 2, 9 and 16 genes, respectively) (**Figure 4A**). Of note, eviction events were mainly found upstream of a small subset of TSSs and appeared transient. This is particularly pronounced in a small group of both primary and secondary response genes (*Ccl3, Ccl4, Rsad2, Tnf, Gbp5, Gbp3, Gbp2, Cxcl10, Oasl1, Tnip3*) that are highly depleted for CpG islands (**Figures 4B** and **Suppl. Fig. 4A**).

**Figure 4.**
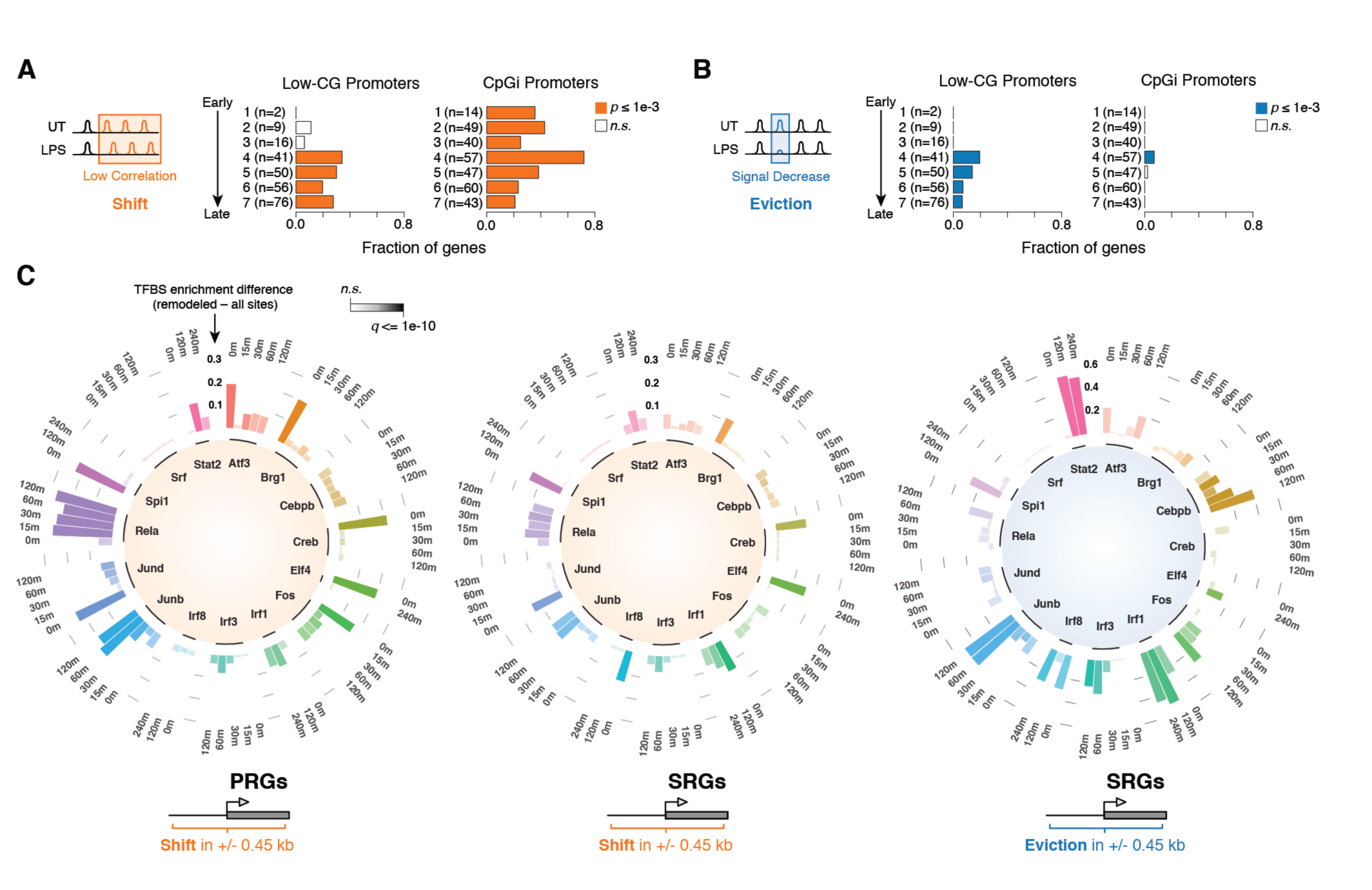
Nucleosome remodeling at different classes of LPS-inducible genes. A-B) Fraction of genes showing nucleosome shift (A) or eviction (B) at their promoter region (± 450 bp), grouped by kinetics of induction upon LPS stimulation (Bhatt et al. 2012) and G+C content. CpGi: CpG island. *P*-values are from a two-sided hypergeometric test. C) Circular, stacked bar plots showing TF binding enrichment at the promoters (TSS ± 450 bp) of LPS-inducible genes exhibiting remodeling events. The analysis is based on ChIP-seq data at the indicated time points of LPS stimulation. Three different groups are shown: shift at primary response genes (PRGs) and at secondary response genes (SRGs), and evictions at SRGs. The *q*-value of the enrichment is color-coded. *p*-values are from a two-sided Chi-squared test, corrected for multiple hypothesis testing using Benjamini-Hochberg.

To determine how TF binding at promoters impacted nucleosome organization, we compiled a collection of TF ChIP-seq profiles (n = 67, including 40 newly generated datasets) encompassing 27 TFs in untreated and LPS-treated macrophages. The quality of the TF ChIP-seq datasets used was assessed by determining the fraction of reads in peaks (FRIP) (**Suppl. Table 1**) and by characterizing the over-represented motifs in each data set (**Suppl. Fig. 5**). In addition to typical stimulus activated transcription factors (NF-kB, AP-1, CREB, IRF, STAT, SRF) we included two myeloid lineage transcription factors (PU.1 and C/EBPβ) (Feng et al. 2008; Goode et al. 2016) and the ETS family member ELF4, which is selectively associated with active *cis*-regulatory elements, contributing to their basal activity (Curina et al. 2017). We also included total RNA Pol II and BRG1, the active subunit of the Swi/Snf chromatin remodeling complex. Basal or LPS-induced TF binding enrichment was measured at the promoters (TSS ± 450 nt) of both primary and secondary response genes showing remodeling (**Figures 4C** and **Suppl. Fig. 4A**, **Suppl. Table 4**). This analysis revealed that: *i*) constitutive binding of PU.1 (SPI1) was associated with all types of remodeling events; *ii*) constitutive binding of ELF4 was associated with nucleosome shifts, likely reflecting the high transcriptional activity of ELF4-associated promoters (Curina et al. 2017); *iii*) basal and inducible IRF1 (Interferon Regulatory Factor 1) binding along with inducible IRF3 and STAT2 binding were selectively associated with remodeling events (especially evictions) at the TSS of secondary response genes, which include genes activated in a paracrine/autocrine manner by early released IFNβ; *iv*) inducible NF-kB binding was associated with shifts but not evictions, and this association was stronger for primary than for secondary response gene promoters. The correlation between basal, inducible and reduced TF binding events and nucleosome remodeling at different classes of genes is highlighted in **Suppl. Fig. 4B**.

Overall, these results indicate that recruitment of different TFs to the promoters of LPS-inducible genes elicit distinct types of nucleosome remodeling events, which in turn correlate with the DNA sequence features of the promoters.

### Nucleosome remodeling at stimulus regulated enhancers

We then investigated nucleosome remodeling at genomic regions showing either constitutive or LPS-induced accessibility. To this end, we generated ATAC-seq profiles at multiple time points of LPS stimulation (30’, 1h, 2h and 4h) (**Suppl. Fig. 6**). While ∼60,000 ATAC-seq peaks were accessible prior to stimulation, the accessibility of 24,252 TSS-distal and 2,471 TSS-proximal peaks was affected by the LPS treatment (**Figures 5A** and **Suppl. Fig. 6A**, **D-G**). ATAC-seq signal gains and losses exhibited broad kinetic complexity, including peaks that were selectively gained or lost at either early or late time points (**Suppl. Fig. 6B-C**). We thus focused on 19,111 TSS-distal, extragenic regions marked by H3K4me1 that were either basally accessible or differentially accessible at one or more time points after LPS stimulation (**Figure 5A**). We then applied our analytical framework (**Figure 3A**) to identify nucleosome remodeling events at these 19,111 putative enhancers. While no significant nucleosome eviction or locally increased occupancy was detected within these regions, 2,135 enhancers (11.2%) exhibited one or more nucleosomal shifts (**Figure 5A** and **Suppl. Table 5**). These events showed a significant enrichment for symmetric sites (*p*-value < 2.2e-16, Chi-squared test). Nucleosomes occupying these remodeled regions were significantly fuzzier (as determined by the DANPOS2 pipeline) (Chen et al. 2013) than nucleosomes at regions where remodeling was not detected (**Figure 5B** and **C**, left panel). These remodeled, accessible regions were instead flanked by more positioned nucleosomes (**Figure 5B** and **C**, right panel). Representative examples of asymmetrically remodeled nucleosomes in the *Ccl5* locus are shown in **Figure 5D**.

**Figure 5.**
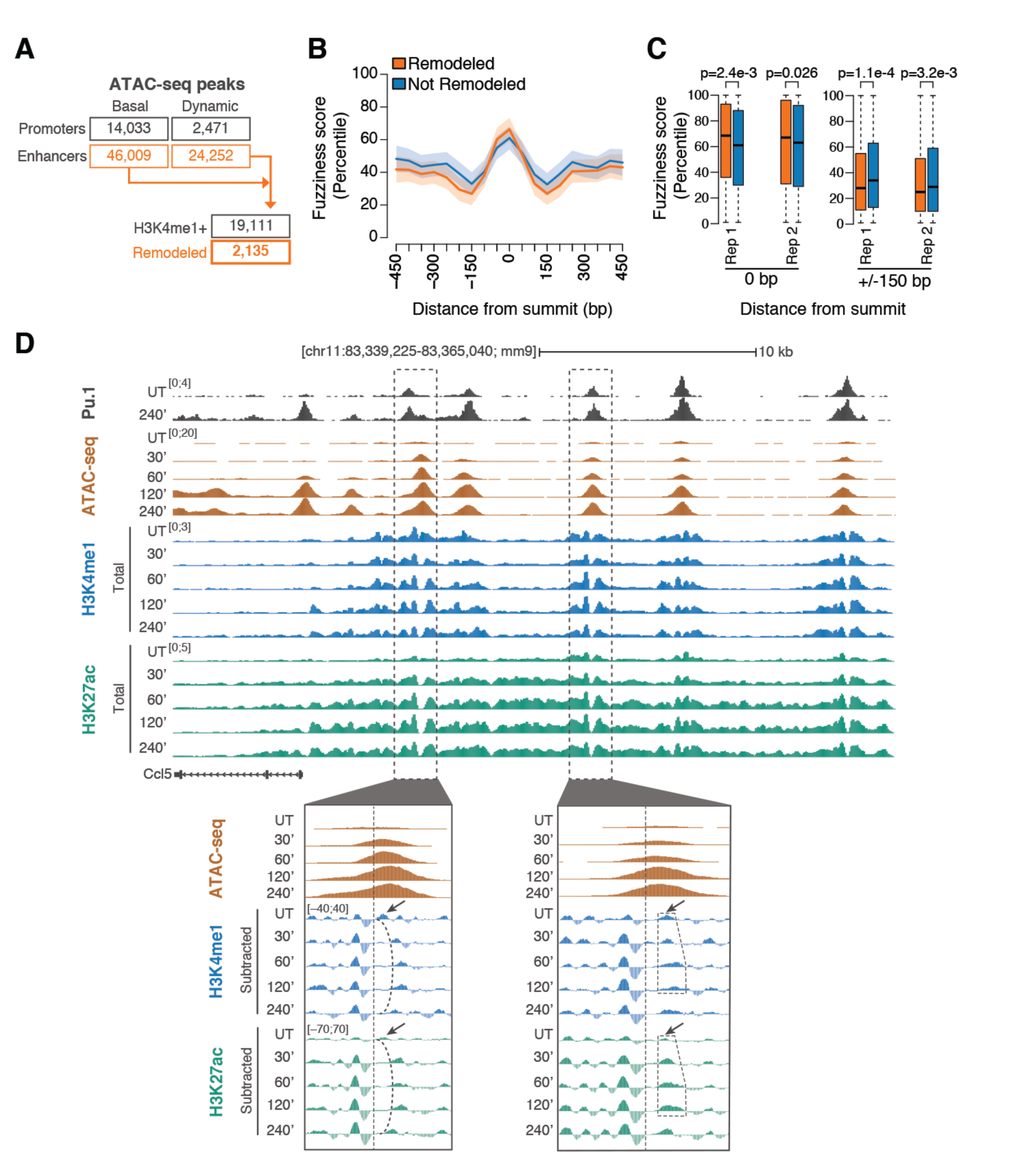
Nucleosome remodeling events at putative enhancers. A) Summary of nucleosome remodeling events at accessible regulatory elements identified by ATAC-seq. Accessible regions either in basal conditions or after LPS stimulation, and either proximal or distal to annotated gene-TSSs, are indicated. B) Nucleosome fuzziness at TSS-distal remodeled (orange) and not remodeled (blue) regions. Data are shown in a window of ± 450 nt centered on the summit of ATAC-seq peaks. Confidence intervals were estimated from a spline fit of the median fuzziness of two biological replicates. C) Distribution of fuzziness scores (percentiles) at the ATAC-seq peak summit (0 bp) and 150 bp upstream or downstream (± 150 bp), for remodeled and not remodeled regions, and each replicate. *p*-values are from a two-sided Wilcoxon rank-sum test. D) Snapshots highlighting representative nucleosome remodeling events at two LPS-inducible accessible regulatory regions upstream of *Ccl5*.

These observations suggest that nucleosomes with intrinsically fuzzier positioning located at enhancer cores are susceptible to remodeling, while those with a stronger positioning are often unperturbed by the landing of TFs in their close proximity.

### Dynamic transcription factor occupancy and nucleosome remodeling events

To investigate the relationship between TF-binding and changes in nucleosome organization at enhancers, we analyzed the panel of TF ChIP-seq profiles described above (**Suppl. Table 1**). We first defined master sets of regulatory regions showing at least one LPS-induced (or LPS-reduced) TF-binding event, at both TSS-distal and TSS-proximal sites (**Suppl. Table 6**). We then correlated the number of distinct TFs that bind each regulatory region and the occurrence of remodeling (**Figure 6A**). We found significant, positive correlation between the number of recruited TFs and changes in nucleosome organization, both at putative enhancers (Spearman’s rank correlation coefficient of 0.74, *p*-value = 4.6e-25) and at promoters (r = 0.66, *p*-value = 1.6e-18). This result suggests that the higher the number of TFs recruited to enhancers, the higher the probability of remodeling, which in turn indicates that remodeling is a tightly regulated, combinatorial process.

**Figure 6.**
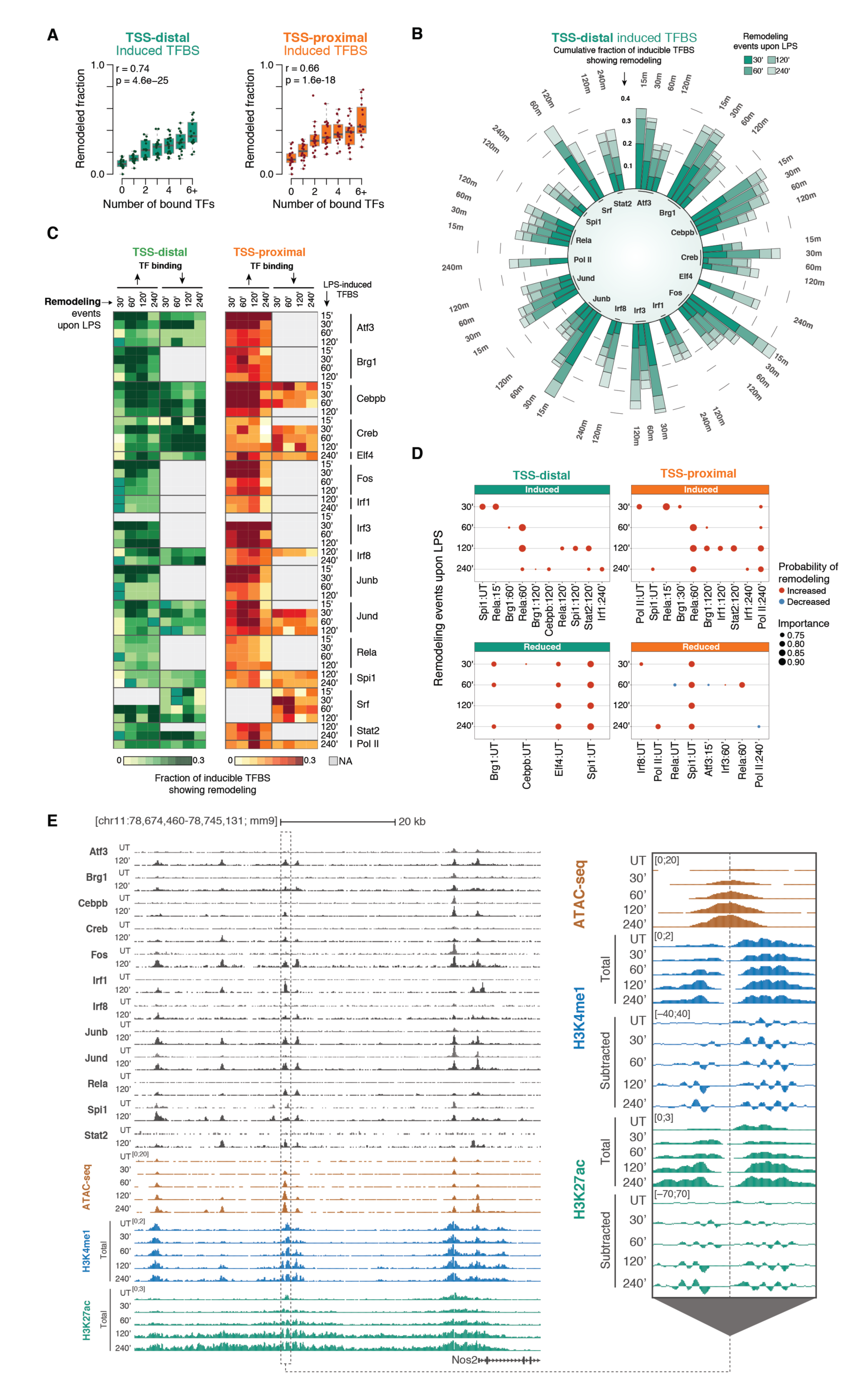
Transcription factor binding and inducible nucleosome remodeling at promoters and enhancers. A) Fraction of putative enhancers (left) and promoters (right) undergoing detectable nucleosome remodeling as a function of the number of inducible TF binding events within a ± 450 nt window. B) Circular stacked bar plots showing the cumulative fraction of inducible TF binding at different time points following LPS stimulation that overlap nucleosome remodeling events. C) Heatmaps summarizing the occurrence of nucleosome remodeling at sites of increased or reduced TF binding at different time points after LPS stimulation. NA: missing data. D) Top-ranked dynamic TF binding events (e.g. Rela:60’ denotes binding of Rela occurring at 60’ after LPS) associated with remodeling events at a given time point after LPS stimulation, selected by bootstrap-lasso logistic regression models. UT: untreated (basal). E) A representative genomic region upstream of *Nos2* is shown. On the left, selected TF ChIP-seq tracks, ATAC-seq tracks and non-subtracted H3K4me1 and H3K27Ac tracks are shown. The inset on the right shows subtracted H3K4me1 and H3K27Ac track at the indicated region.

Next, for each TF we measured the fraction of LPS-inducible changes in TF binding (induction or reduction) that was associated with detectable nucleosome remodeling at the TF recruitment site at different time points. For the majority of TFs, most remodeling events occurred within 60’ of LPS, with additional later events becoming progressively less common over time (**Suppl. Fig. 6B-C** and **7**, and **Suppl. Table 7**). This is consistent with the genome-wide pattern of LPS-inducible accessibility at putative regulatory elements (**Suppl. Fig. 6D** and **6F**). A fine-grained dissection of remodeling events by time points (**Figure 6C**) revealed that while these occurred throughout the entire stimulation kinetics, they became less common at the last time point analyzed (240’), which is likely a reflection of the overall reduction in signaling strength at later time points after LPS stimulation. Importantly, the analysis of ChIP-seq profiles at the early time points (15-30’ after LPS) indicates that in many cases (e.g. IRF3, RELA, JUNB) remodeling events associated with early TF recruitment persisted over time.

Finally, we set out to test whether TF dynamics could jointly predict remodeling events. To this end, we trained lasso logistic regression models at distal and proximal sites and estimated the relative importance of each TF with respect to remodeling events at a given time point. We found that model performances were generally modest but significantly higher at distal sites (AUC = 0.64-0.70) than at proximal sites (AUC = 0.58-0.64). Feature importance analysis identified TF binding events associated with increased or decreased probability of occurrence of a remodeling event. LPS-inducible binding of NF-kB (RELA) within 60’ of stimulation was the most single predictive feature of remodeling (selected >90% of the times) at both TSS-distal and proximal sites. Moreover, RELA recruitment at 60’ predicted remodeling also at later time points, indicating that nucleosome alterations induced by NF-kB are sustained over time. Late (120’-240’) binding of IRF1 and STAT2 was also moderately associated to remodeling (selected >75% of the times) at these regions. In contrast, PU.1 (SPI1) binding in resting conditions was strongly associated (selected >90% of the times) to remodeled regions showing reduced TF-binding upon LPS stimulation (**Figure 6D**). The snapshot in **Figure 6E** shows the chromatin changes and TF recruitment at regulatory regions upstream of the LPS-inducible gene *Nos2*. The region highlighted in the inset shows a nucleosome remodeling event identified by our pipeline and associated with a gain in the ATAC-seq signal. The H3K4me1-marked nucleosomes on both side showed increased positioning after remodeling of the central fuzzy nucleosome.

Overall, these data provide a comprehensive overview of the impact of signal-induced recruitment of individual TF on nucleosome remodeling. They show that only a fraction of TF recruitment events is associated with detectable nucleosome remodeling and indicate that individual TFs, whereas acting combinatorially to displace nucleosomes, appear to have different remodeling efficiencies.

## DISCUSSION

In mammalian cells, nucleosome organization at *cis*-regulatory elements is determined by a complex interplay between transcription factors bound to chromatin and DNA sequence features that either facilitate or disfavor nucleosome assembly (Tillo et al. 2010; Valouev et al. 2011; Barozzi et al. 2014; Lai and Pugh 2017). A general paradigm is that the unique nucleosome profile characteristic of a given cell type is largely driven by two opposing forces: on the one hand, enhancers have an intrinsically high propensity to assemble nucleosomes, thus generally making their regulatory information inaccessible unless appropriate combinations of TFs are expressed (Tillo et al. 2010; Barozzi et al. 2014); on the other hand, lineage-determining TFs actively maintain a fraction of the genomic regulatory repertoire in a nucleosome-depleted state, regulating its accessibility in a cell type-specific manner (Barozzi et al. 2014). The barrier effect generated by lineage-determining TFs bound to enhancers represents a major force driving the precise positioning of nucleosomes at the two sides of accessible enhancer cores.

In this study, we set out to investigate a critical and unaddressed issue in this area, namely the interplay between pre-existing nucleosome organization and the recruitment to *cis*-regulatory elements of TFs activated by acute stimulation. A widely-accepted model is that TFs activated by stimulation land at enhancers that are pre-marked and thus kept accessible by the lineage-determining TFs characteristic of a given cell type (Glass and Natoli 2016). In this way, transcriptional responses to otherwise identical stimuli are contextualized and rendered cell type-specific. In this model, binding of stimulus-activated TFs is highly opportunistic and in fact restricted to genomic regions constitutively bound by lineage-determining TFs, which displace nucleosomes and maintain the underlying DNA recognition motifs exposed and accessible. A corollary of this model is that since inducible binding of stimulus-activated TFs occurs within accessible regions, it should not significantly impact nucleosome depletion. However, this inference is not easily reconciled with the observation that acute stimulation also increases accessibility of a subset of enhancers, as determined by techniques such as ATAC-seq or DNAseI-seq (Novakovic et al. 2016; Park et al. 2017). The nature of chromatin alterations leading to such changes in accessibility induced by stimulation is unknown. In particular, it is unclear if the recruitment of stimulus-activated TFs suffices to displace well-positioned nucleosomes in order to promote a further expansion of the accessible repertoire of regulatory elements. Our data indicate that nucleosomes undergoing remodeling in response to stimulation are significantly less positioned than non-remodeled nucleosomes. These fuzzy nucleosomes are frequently flanked by well-positioned nucleosomes that are refractory to the nucleosome remodeling activity of stimulus-inducible TFs recruited in their immediate vicinity.

These data support a model in which stimulus-responsive *cis*-regulatory elements are frequently flanked by immobile, well-positioned nucleosomes, while their core is wrapped into intrinsically mobile and/or partially accessible nucleosomes amenable to rapid displacement or reorganization upon inducible recruitment of TFs. Therefore, the overall nucleosomal organization of stimulus-responsive enhancers is extremely robust to extrinsic perturbations, a feature that might prevent deviations from the differentiated state when cells are exposed to perturbations. Such resilience of the nucleosomal landscape likely reflects the dominance of the nucleosome-organizing activity of lineage-determining TFs such as PU.1 (Barozzi et al. 2014), whose overall genomic distribution is only marginally affected by stimulation (Ostuni et al. 2013; Mancino et al. 2015).

Our study also revealed that co-binding of multiple TFs to *cis*-regulatory elements is tightly associated with remodeling. This result suggests that even remodeling of fuzzy nucleosomes might be a highly cooperative process in which multiple collaborating transcription factors provide an important contribution. Nevertheless, not every TF appears to elicit remodeling with the same efficiency. Indeed, some TFs bound with a similar frequency to remodeled and unaffected nucleosomes, whereas others were strongly associated with remodeling events. The most unexpected finding in this regard relates to NF-kB, whose DNA-binding domain is structurally incapable of accommodate nucleosomal DNA (Natoli et al. 2005; Lone et al. 2013). Nevertheless, when recruited to DNA, NF-kB appeared to strongly associate with the disruption of local nucleosome organization, consistently predicting nucleosome remodeling after LPS stimulation. As NF-kB plays a pivotal role in the induction of the inflammatory gene expression program, its association with enhanced chromatin accessibility suggests that this may also depend on its ability to control a feed-forward mechanism acting at promoters and enhancers, whereby NF-kB binding augments local chromatin accessibility thus favoring the subsequent recruitment of additional transcription factors.

Overall, the data reported in this study contribute to clarify how the exposure of mammalian cells to external stimuli impacts the accessible landscape *of cis*-regulatory elements by remodeling mainly poorly positioned nucleosomes. On the other hand, the remarkable resilience of the overall nucleosomal landscape in the face of even very strong stimuli lends support to the notion that maintenance of the nucleosomal organization in a changing environment represents an essential feature of cell differentiation.

## MATERIALS AND METHODS

### *Cell cultures*. Macrophages were derived from bone marrows of C57/BL6 mice (Harlan) as described (Curina et al. 2017)

#### MNase-ChIP-seq

2×10^7^ macrophages were fixed with 1% HCOH, quenched with 0.125 mM Tris and washed twice with cold PBS. Cell pellets were resuspended in a 15 mM NaCl, 15 mM Tris-HCl [pH 7.6], 60 mM KCl, 2 mM EDTA, 0.5 mM EGTA, 0.3 M sucrose buffer (0.5 mM PMSF, 1 mM DTT, 0.2 mM spermine, 1 mM spermidine) and lysed upon addition of 0.4% NP40. Nuclei were washed with a 15 mM NaCl, 15 mM Tris-HCl [pH 7.6], 60 mM KCl, 0.3 M sucrose buffer (0.5 mM PMSF, 1 mM DTT, 0.2 mM spermine, 1 mM spermidine). Digestion was performed with 6 units of MNase (Roche 10107921001) in 500 μl of a 20 mM Tris-HCl [pH 7.6], 5 mM CaCl2 digestion buffer, for 2 hours at 37°C to have around 80% mononucleosomes and 20% di-nucleosomes. The reaction was stopped by adding EDTA to a 50mM final concentration. Digestion was checked on an agarose gel, after decrosslinking of a small fraction of the lysate. The samples were diluted in 3 ml of a 10mM Tris-HCl[pH 8], 100 mM NaCl, 1mM EDTA, 0.5mM EGTA0.1% Na-Deoxycholate, 0.5% Na-Laurylsarcosine buffer, gently sonicated to favor nuclei lysis and then divided into 4 aliquots (corresponding to 5 millions each) for immunoprecipitation with 5 μg of antibody. Antibodies were pre-bound to G protein-coupled paramagnetic beads (Dynabeads) in PBS-0.5% BSA and incubated with lysates overnight at 4°C. Beads were washed six times in a modified RIPA buffer (50 mM HEPES [pH 7.6], 500 mM LiCl, 1 mM EDTA, 1% NP-40, 0.7% Na-deoxycholate) and once in TE containing 50 mM NaCl. DNA was eluted in TE-2% SDS and crosslinks reversed by incubation overnight at 65°C. DNA was then purified by MinElute PCR purification kit (Qiagen) and quantified with PicoGreen (Invitrogen). ChIP DNA was prepared for HiSeq2000 sequencing following standard protocols. The antibodies used included anti H3K4me1 (Abcam Ab 88959), anti-H3K4me3 (Active Motif 39159), anti-H3K27Ac (Abcam Ab4729).

#### ATAC-seq

The original ATACseq protocol (Buenrostro et al. 2013a)(Buenrostro et al., 2013) was modified according to Lara-Astiaso et al. (Lara-Astiaso et al. 2014). Briefly, 5×10^4^ cells were lysed in100 μl of a 10mM Tris-HCl [pH 7.4], 10mM NaCl, 3mM MgCl2, 0.1% Igepal CA-630. Nuclei were pelleted by centrifugation for 20 min at 500g, 4 °C. Nuclear pellet was re-suspended in 25 µl of a 10mM Tris-HCl [pH8.4] and 5mM MgCl2 buffer containing 1 µl of Tn5 transposase (made in house). The reaction was incubated at 37°C for one hour and then stopped by adding 5 µl of clean up buffer (900mM NaCl, 300mM EDTA), 2ul of 5% SDS and 2 µl of Proteinase K (20 µg/µl) and incubating the reaction for 30 min at 40 °C. Tagmented DNA was purified using 2× SPRI beads (Agencourt AMPure XP-Beckman Coulter). Finally, 2 µL of 10 µM indexing primers and KAPA HiFi HotStart ReadyMix were used to PCR-amplify the obtained DNA library of tagmented DNA. Fragments smaller than 600 bp were isolated by size selection (using 0.65× SPRI beads) and then purified with 1.8× SPRI beads. ATAC-seq library size was assessed using a Tapestation D5000 High Sensitivity ScreenTape (Agilent). Libraries were sequenced on an Illumina NextSeq 500.

#### Chromatin-associated RNA

10×10^6^ cells for each time point were lysed with an ice cold 10 mM Tris-HCl [pH 7.5], 0.05% NP40, 150 mM NaCl buffer for 5 min. The lysate was then layered on 2.5 volumes of a chilled sucrose cushion (24% sucrose in lysis buffer) and centrifuged for 10 min, 4°C, 14,000 rpm. The nuclei pellet was gently rinsed with ice-cold PBS/1 mM EDTA, then resuspended in 20 mM Tris-HCl [pH 7.9], 75 mM NaCl, 0.5 mM EDTA, 0.85 mM DTT, 0.125 mM PMSF, 50% glycerol buffer, by gentle flicking of the tube. An equal volume of cold 10 mM HEPES [pH 7.6], 1 mM DTT, 7.5 mM MgCl, 0.2 mM EDTA, M NaCl, 1 M UREA, 1% NP-40 buffer was added. The tube was gently vortexed and then centrifuged for 2 min, 4°C,14,000 rpm. The chromatin pellet was rinsed with cold PBS/1 mM EDTA and then dissolved in TRIzol (Invitrogen). Chromatin-associated RNA was purified with TRIzol, with an additional phenol/chloroform extraction step prior to precipitation. Purified RNA was used for library preparation with the Illumina Truseq V2 protocol. Libraries were sequenced on an Illumina HiSeq2000 (50-bp single reads).

#### Gene, promoters and enhancers annotations

Models for UCSC known gene were downloaded from iGenome (https://support.illumina.com/sequencing/sequencing_software/igenome.html) on July 17, 2015. Unless otherwise stated, a TSS-proximal region was defined as ± 2.5 kb from a TSS. Enhancers were defined as TSS-distal, Pu.1-bound, H3K4me1-positive regions (Barozzi et al. 2014). Regions on chrM were excluded.

#### MNase-ChIP-seq data analysis

Reads were aligned to the reference genome (mm9) using bowtie (v0.12.7) (Langmead et al. 2009) with parameters -v 2 -m 1, ensuring only unique-mapping aligned reads with two or fewer mismatches were retained. Enriched regions were identified using MACS v1.4 (Zhang et al. 2008) with parameters --nomodel --nolambda --pvalue=1e-5 --bw=100 --shiftsize=75 --gsize=mm. Wiggle tracks for visualization on the UCSC genome browser (Kent et al. 2002) were generated using MACS v1.4 and rescaled to RPM (Reads Per Million sequenced reads; de-duplicated reads). SAMtools (v0.1.19) (Li et al. 2009) was used to split the reads by strand, then *genomeCoverageBed* from BEDTools (v2.19.1)(Quinlan and Hall 2010) was used to generate the strand-specific, genome-wide coverage profiles at single base-pair resolution. The minus profile was then subtracted from the plus profile using *wigmath.subtract* from Java Genomics Toolkit (https://github.com/timpalpant/java-genomics-toolkit). For comparative analyses, the derivative of the cumulative signal was calculated from previously published MNase-seq data in untreated macrophages (pool of four biological replicates)(Barozzi et al. 2014). SAMtools (*samtools view -s*) was used to subsample the total reads.

#### ATAC-seq data analysis

Reads were aligned to the reference genome as described for the MNase-ChIP-seq profiles. Accessible regions were then identified using MACS v1.4 (Zhang et al. 2008) with parameters --gsize=mm --bw=300 --nomodel --nolambda --shiftsize=150. Wiggle profiles were also generated as described for the MNase-ChIP-seq profiles. First, accessible regions from each sample were split into TSS-distal and TSS-proximal as described above. Master lists of peaks were generated merging all the replicates and all time points, but separating TSS-distal and TSS-proximal, using *mergeBed* from BEDTools. A region was considered for further analyses if showing a peak in 2 or more replicates, in at least one time point. Raw read counts over these regions were quantified using *coverageBed* from BEDTools. *edgeR* (v3.20.9) was then used for normalization and for estimating differentially accessible regions between each time point and the untreated. *estimateCommonDisp* and *estimateTagwiseDisp* were run separately, followed by Trimmed Mean of M-values (TMM) normalization. Differential accessibility was evaluated by *exactTest*, followed by a filter on an absolute fold-change > = 2 and a *q*-value < = 0.05 (Benjamini-Hochberg correction)(Benjamini and Hochberg 1995). A master lists of intergenic, accessible regions was derived from the TSS-distal intervals. Regions showing accessibility in the untreated (either changed or unchanged by LPS) or gaining accessibility upon LPS treatment were considered. A filter was applied to keep only those regions with consistent H3K4me1 between replicates (untreated). This filter ensures a fair estimation of nucleosome remodeling across all the regions, whether constitutively active or induced by LPS.

#### TF-ChIP-seq data analysis

FASTQ files were downloaded from the Gene Expression Omnibus (GEO)(Barrett et al. 2013), and processed using Trim Galore (−q 20; https://www.bioinformatics.babraham.ac.uk/projects/trim_galore/). Reads were aligned to the reference genome as described for the MNase-ChIP-seq profiles. TF-bound locations were identified using MACS v1.4 (Zhang et al. 2008) with parameters --gsize=mm --bw=300 --nomodel --shiftsize=100, and sample-matched input DNA as control. LPS-induced and reduced events were identified using the same parameters. Those locations showing differential binding and overlapping a significant peak versus input DNA were kept. Unless specified otherwise, only peaks showing a *p*-value < = 1e-10 were retained for further analyses. Master lists of TF-binding events (i.e. regions bound by at least one TF, at any time point), were derived separately for LPS-induced and reduced events, independently for TSS-proximal and TSS-distal elements, as follows. First, summits within 450 bp from each other were clustered together using *mergeBed* from BEDTools. A summit for each cluster was then calculated as the average of all the summit positions of the peaks included in the cluster.

#### Chromatin-associated-RNA-seq data analysis

Reads were aligned to the indexed murine cDNA (assembly GRCm38, release 92) using Kallisto *quant* (v0.44.0) (Bray et al. 2016), with parameters -b 100 -l 300 -s 100 --single. Sleuth (R package v0.29.0)(Pimentel et al. 2017) was used to identify differentially expressed genes (DEGs) upon LPS stimulation. Sleuth identifies DEGs by comparing a full to a reduced model. Here, the reduced model considered only replicates while a full model replicates and time of stimulation. Both models were fit using *sleuth_fit* and compared using *sleuth_lrt*. DEGs were defined if *q*-value < = 0.05 and an absolute 2-fold-change. For LPS-induction, genes up-regulated as early as 30’ or 60’ were considered early-response genes, as opposed to late-response genes, i.e those showing increased expression only at 120’ or 240’.

Genes were classified as either primary- or secondary-response genes (PRG or SRG) using published gene expression profiles from stimulated macrophages in the presence of cycloheximide (CHX) and Type I IFN receptor (IFNAR)-deficient (Ifnar^−/−^) stimulated macrophages (Tong et al. 2016). Given a time point, LPS-induced genes down-regulated by ≥ 70% in either CHX-treated or Ifnar^−/−^ macrophages, were considered as SRGs. Genes that were not defined as SRG at any time point were classified as PRGs.

#### Characterization of symmetric and asymmetric nucleosomal arrays at putative enhancers

The orientation of nucleosomal patterns was inferred by the ratio of normalized signals on the two sides of the Pu.1 summit. These coverages were calculated using *coverageBed* from BEDTools. Spatially resolved signals were extracted for: (1) H3K4me3 and total RNA polymerase II ChIP-seq profiles in untreated macrophages (Ostuni et al. 2013); (2) CTCF (ChIP-seq profile in untreated macrophages; Ghisletti & Natoli, unpublished); (3) the PAS (PolyA Signal) using a model as described (Austenaa et al. 2015).

#### Statistical learning

Least absolute shrinkage and selection operator (lasso) logistic regression models were used to discriminate symmetric from asymmetric nucleosomal patterns at intergenic enhancers based on DNA sequence features, and to predict nucleosome remodeling events associated with TF dynamics during LPS stimulation. Models were trained and evaluated as described (Comoglio et al. 2015; Comoglio et al. 2018).

For modeling of nucleosomal symmetry patterns at enhancers, DNA sequences were extracted from the reference genome (mm9) are scored as follow. Three sets of features were considered:

1. DNA sequence content encoded as *k*-mers (2 ≤ *k* ≤ 4), within 450 bp up- and down-stream of the PU.1 summit.

2. Average DNA shape feature values within the same window, computed using the R package *DNAshapeR* v1.5.3 (Chiu et al. 2017). Thirteen DNA shape features were used in this study (Sagendorf et al. 2017), along with predicted minor-groove electrostatic potential (Chiu et al. 2017).

3. Transformed FIMO *p*-values (−10*log10(p)) (Grant et al. 2011) for a curated collection of >1,700 position weight matrices (PWMs) representing mammalian TF motifs (Diaferia et al. 2016), core promoter elements, 5’ splice site and polyA signal motifs. These were computed within 300 bp of the PU.1 summit as previously described (Barozzi et al. 2014) using FIMO from MEME v4.11.3. To train the classifiers, data points were randomly partitioned into 10 balanced training (80%) and test (20%) sets. Lasso logistic regression models were trained in cross validation (ten-fold) using the R package glmnet v2.0-13 (Friedman et al. 2010). The value of the regularization parameter resulting in a cross-validated misclassification error within one standard error from the minimum was used to predict the class labels of the enhancers in the test set. Model performances were evaluated by computing the area under the receiver operating characteristic curve (AUC) using the R package *ROCR* v1.0-7(Sing et al. 2005) and average AUC values were computed. Feature importance analysis was performed via bootstrap-lasso as previously described (Comoglio and Paro 2014). Features with selection probability (importance) ≥ 0.85 were considered.

The same approach was used to model nucleosome remodeling events associated with TF dynamics, using TF ChIP-seq signals (RPKM) at dynamic peaks as input features.

#### Detection and classification of nucleosome remodeling events and downstream analyses

Analyses were run on pre-defined set of regions, based on the reproducibility of either H3K4me3 (for TSS-proximal regions) or H3K4me1 (for TSS-distal regions) patterns in the untreated condition and in the LPS time point under consideration. These sets considered the peak intersection of the two biological replicates. Local losses (eviction) or gains in MNase-ChIP-seq signals upon LPS stimulation were identified as local significant changes within globally unaffected broader regions as follows. First, the differences between LPS and untreated were computed using a fixed window of 5.4 kb (450 bp * 12, defining the broad context). Second, each region was scanned using a sliding window of 450 bp (defining the local context), with a step of 150 bp. Sliding windows exhibiting a statistically significant difference occurring within a fixed window displaying no significant difference were classified into losses or gains based on sign. Formally, the difference for each window was assessed using *csaw* (R package v1.12.0)(Lun and Smyth 2016). After normalization and estimation of dispersion, a quasi-likelihood negative binomial generalized log-linear model was used to fit the count data and to estimate the significance of differential signals. A *q*-value < = 0.05 (Benjamini-Hochberg correction)(Benjamini and Hochberg 1995) was applied. To identify nucleosomal shifts upon LPS stimulation, reproducible decreases in signal correlation between two conditions were detected using strand-specific subtracted MNase-ChIP-seq coverage profiles. A sliding window (450 bp in size, with a step size of 150 bp) was used to scan a 2.25 kb region centered on the landmark of interest (i.e. the TSS of a gene or the summit of a peak). For each window, the subtracted MNase-ChIP-seq coverage was extracted and individually summarized at 15 bp resolution (*n* = 30 bins) for each replicate and condition. Then, the Spearman’s rank correlation coefficient of the summarized coverage between the two untreated samples and each of the four possible untreated-LPS treated sample pairs was computed. We then determined whether the correlation between the summarized coverage of a given untreated replicate and a given LPS-treated replicate was significantly smaller than the correlation between the two untreated replicates, for each of the four possible untreated-LPS treated pairs. The statistical significance of the difference between these dependent correlations was determined using the *paired.r* function from the R package *psych*. The maximum *p*-values from these four tests was used as conservative estimate of significance. *P*-values from the same 2.25 kb region were combined using the Simes’ method (Sarkar and Chang 1997), and further corrected for multiple hypothesis testing by Independent Hypothesis Weighting (*IHW*; R package v1.6.0)(Ignatiadis et al. 2016). The geometric mean of the signal coverage of the four samples (two untreated and two LPS-treated) was used as covariate.

Nucleosome fuzziness was assessed genome-wide using the function *dpos* of DANPOS (v2.2.2)(Chen et al. 2013) with default parameters. Only those estimates overlapping a peak called from the same MNase-ChIP-seq profiles were considered. The fuzziness scores were then converted to percentiles.

#### Data and software availability

Raw sequencing data was deposited at the Gene Expression Omnibus (GEO) under accession number GSE119693.

## ACKNOWLEDGEMENTS

This study was supported by the European Research Council (ERC AdG #692789 to GN). F.C. was supported by an EMBO long-term fellowship (1305-2015 and Marie Curie Actions LTFCOFUND2013/GA-2013-609409) and a Swiss National Science Foundation postdoctoral fellowship (P2EZP3_165206). I.B. was funded through an Imperial College Research Fellowship. We thank Boris Lenhard (Imperial College London) for discussions; Luca Rotta, Thelma Capra and Salvatore Bianchi (IEO and IIT Center for Genomic Sciences, Milan) for the preparation and processing of part of the NGS libraries. F.C. would like to thank Tony Green and Bas van Steensel for supporting this project.

## AUTHOR CONTRIBUTIONS

Data generation: MS, SP and XL. Data analysis and modeling: FC and IB. Funding acquisition and supervision: GN and STS. All authors contributed to writing, reviewing, and editing.

## SUPPLEMENTAL FIGURES

**Supplemental Figure 1.**
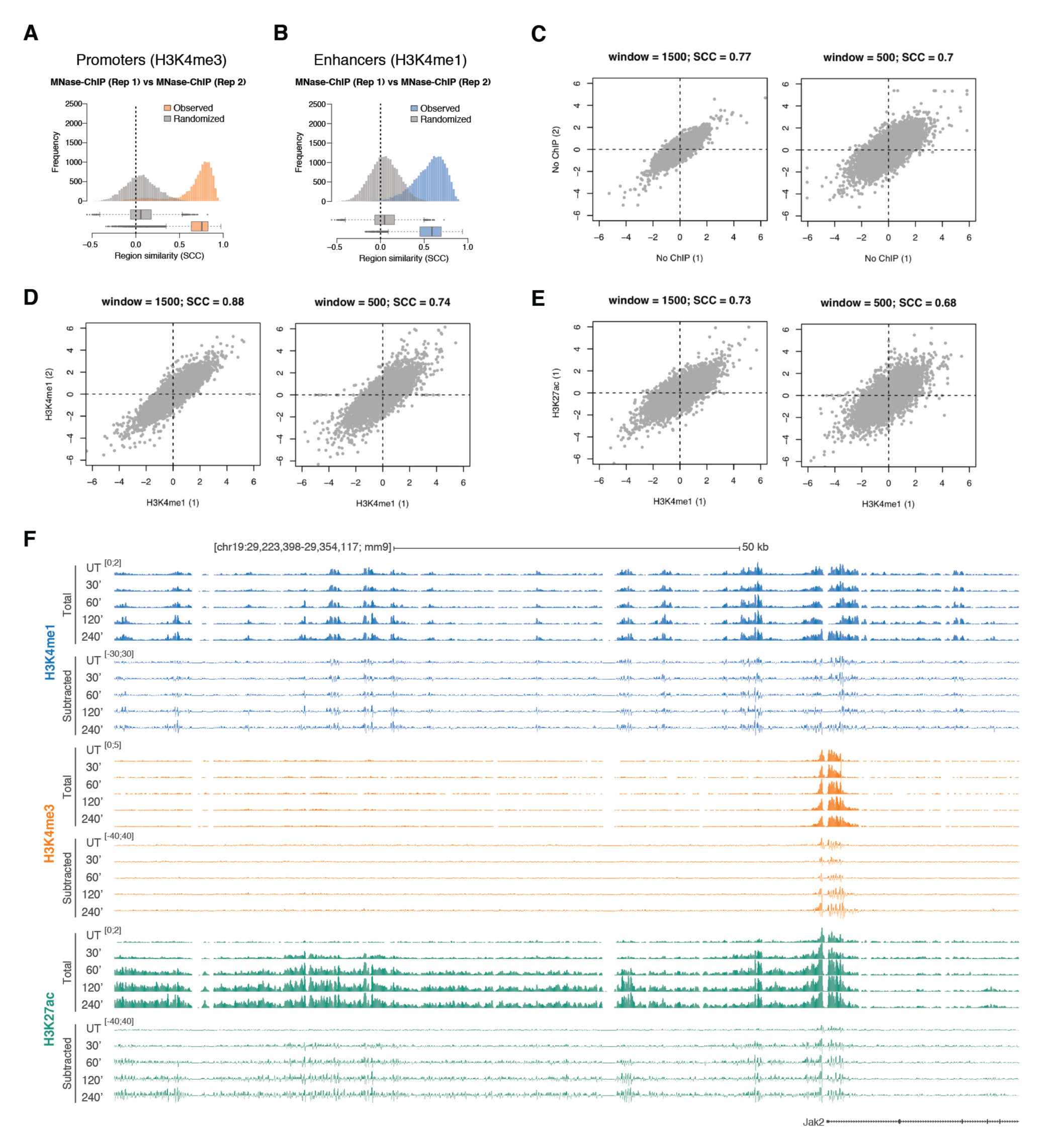
Reproducibility of MNase-ChIP-seq data. A) Distribution of Spearman’s rank correlation coefficient (SCC) between biological replicates of H3K4me3 MNase-ChIP-seq signals within promoters. Distributions for randomized sets are shown in grey. B) Same as (A), for H3K4me1 MNase-ChIP-seq signals at enhancers. C-E) Spearman’s rank correlation coefficient (SCC) between ratios of nucleosomal signals (upstream/downstream) at enhancer cores across replicates of MNase-seq (top), H3K4me1 MNase-ChIP-seq (middle) and H3K27ac MNase-ChIP-seq (bottom). Results for two window sizes (± 1.5 kbp and 0.5 kbp) are shown. F) A representative snapshot of an extended (130 kb) genomic region.

**Supplemental Figure 2.**
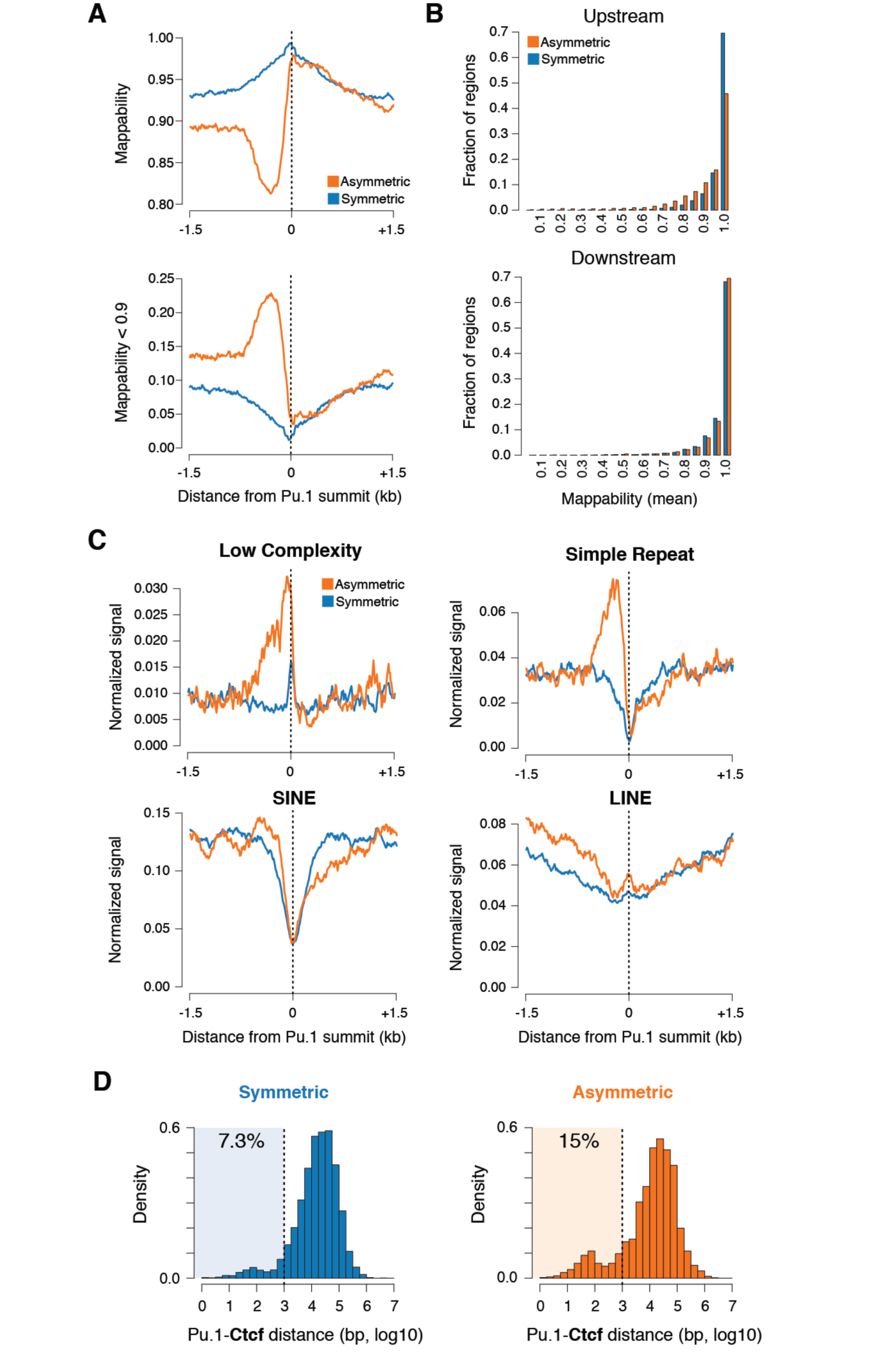
Symmetric and asymmetric enhancers: reproducibility and impact of mappability and repeats. A) Distribution of average mappability (top) and fraction of regions showing mappability below 0.9 (for the considered bin; bottom). Asymmetric and symmetric sites shown in orange and blue, respectively (bin size = 10 bp). B) Distribution of average mappability within upstream and downstream genomic regions. C) Distribution of the indicated repetitive elements and low-complexity regions upstream and downstream of the cores of asymmetric and symmetric enhancers. D) Distribution of distances between the summit of a PU.1 peak and the nearest CTCF peak at symmetric and asymmetric enhancers.

**Supplemental Figure 3.**
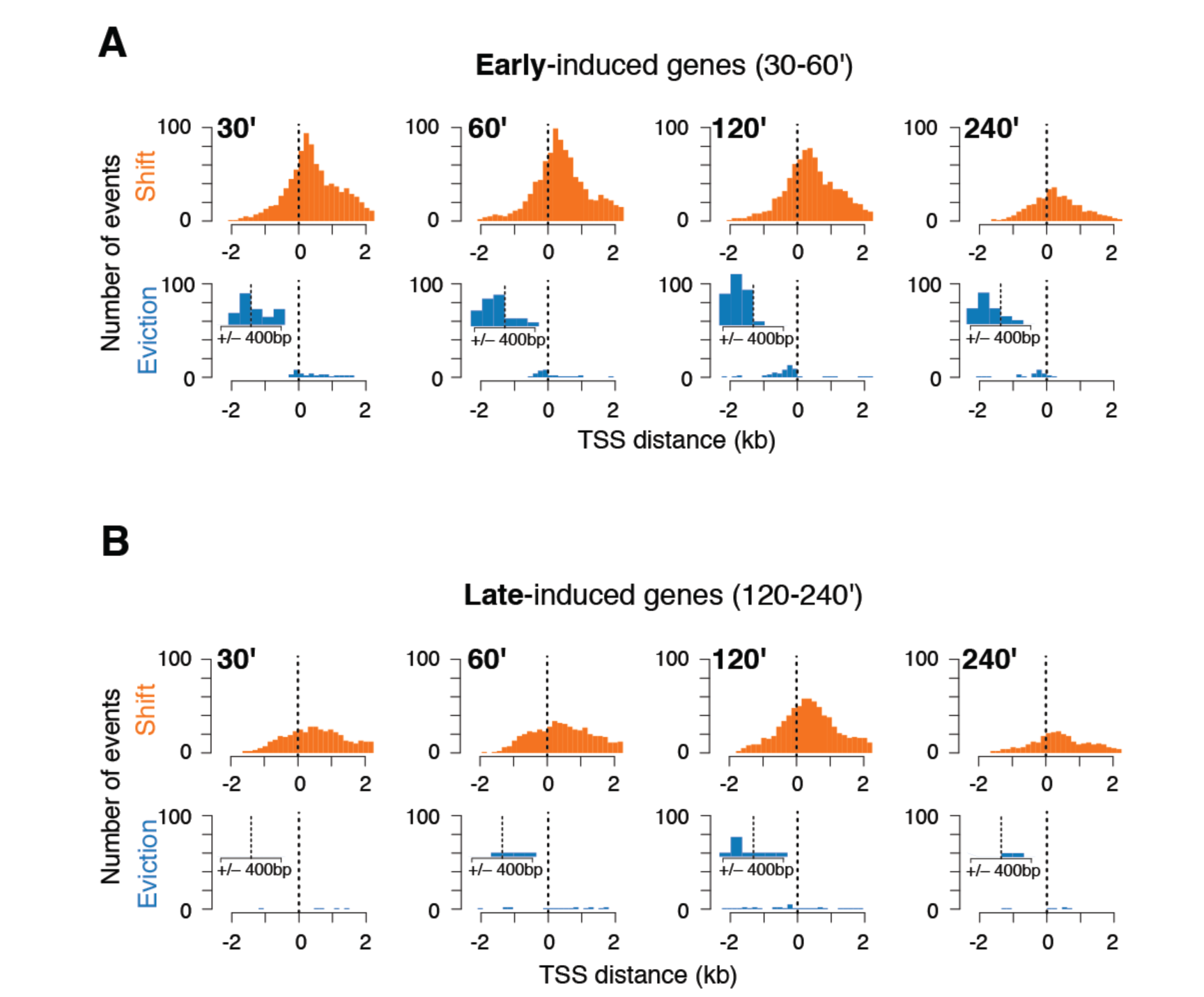
Remodeling events detected at early (30-60’) and late (120-240’) LPS-induced genes. A-B) Distributions of nucleosome remodeling events centered on the TSS of early and late response genes at different time points after LPS stimulation. Insets show an enlargement of the distributions of eviction events throughout the central 900 bps around the TSS.

**Supplemental Figure 4.**
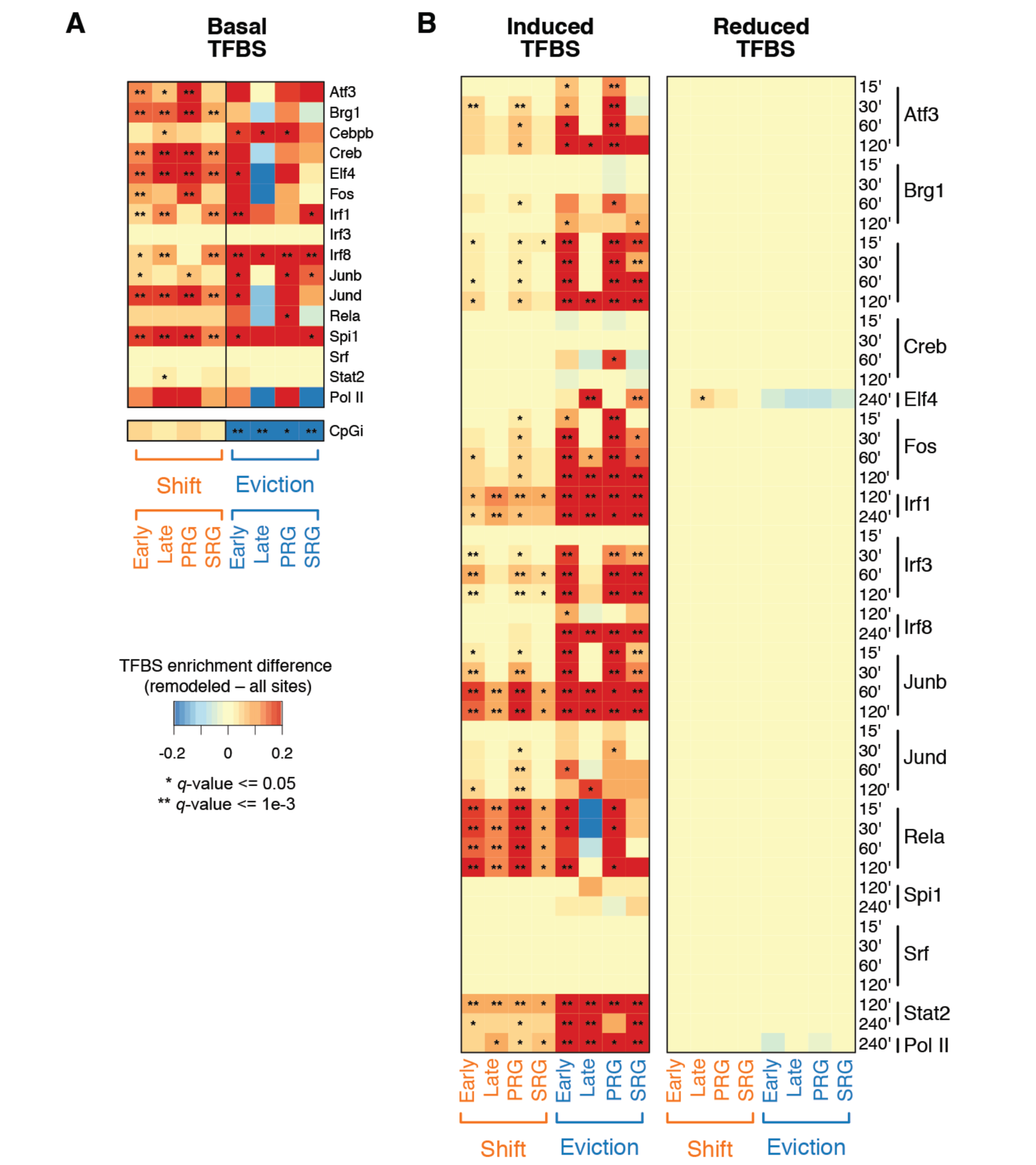
Transcription factor binding and nucleosome remodeling events at LPS-inducible gene promoters. A) Observed-to-expected differences in basal TF binding events (from intersection with ChIP-seq data) at the promoters of LPS-induced genes showing either nucleosome shifts or evictions. Genes were grouped in either early vs late response genes, or primary vs secondary response genes (PRGs and SRGs, respectively). B) Same as (A) but based on LPS-induced and reduced TF-binding events.

**Supplemental Figure 5.**
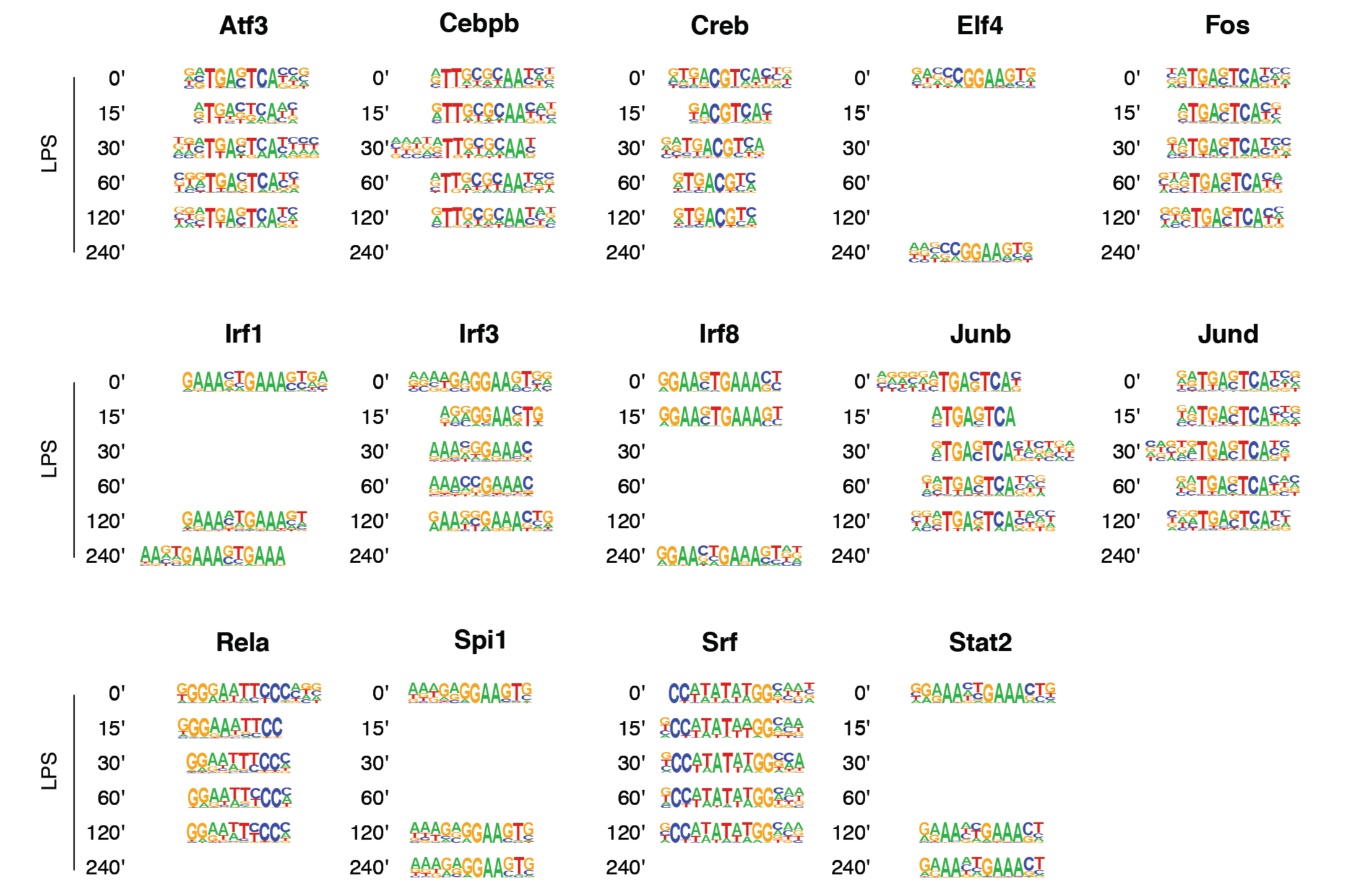
De novo motif discovery analysis on the 1,000 most enriched regions for each TF-ChIP-used in this study.

**Supplemental Figure 6.**
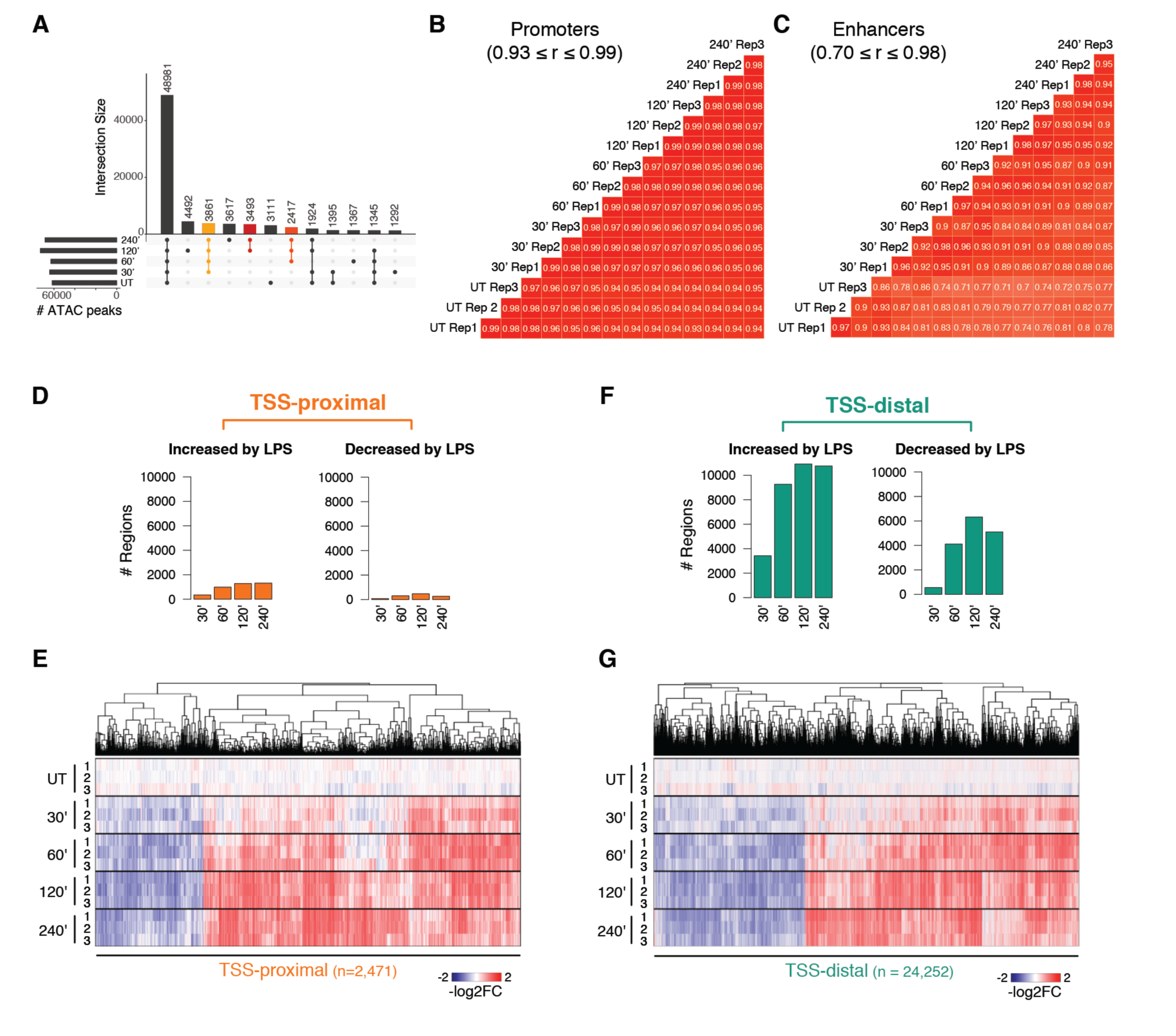
ATAC-seq analysis of chromatin accessibility in LPS-stimulated macrophages. A) Overlap between ATAC-seq peaks identified at the indicated time points during LPS stimulation. B) Spearman’s rank correlation coefficient (SCC) of ATAC-seq signals at promoters, for three biological replicates at the indicated time points during LPS stimulation. Signals were quantified within a 5kb window centered on the TSS. C) Same as (B) but at putative enhancers. D) Total number of TSS-proximal regions whose accessibility was modulated by LPS stimulation. E) Hierarchical clustering of ATAC-seq signals within TSS-proximal regions overlapping at least one ATAC-seq peak during the course of LPS stimulation. Biological triplicates are shown. F-G) Same as (D-E) but for TSS-distal regions.

**Supplemental Figure 7.**
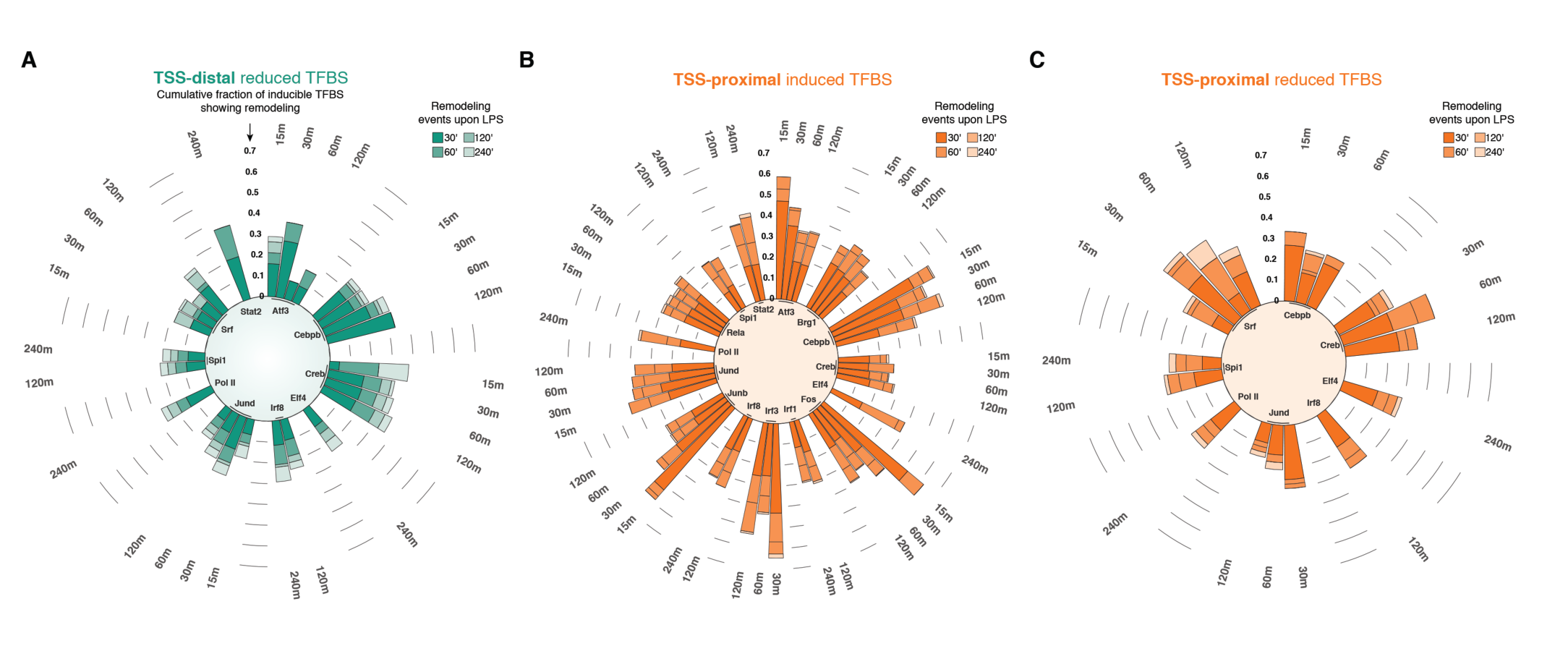
Transcription factor binding and nucleosome remodeling events. A) Circular stacked bar plots showing the cumulative fraction of TSS-distal sites overlapping nucleosome remodeling events and exhibiting reduced TF binding at different time points following LPS stimulation. B-C) Same as (A) but for TSS-proximal sites either induced (B) or reduced (C) by LPS stimulation.

## SUPPLEMENTAL TABLE LEGENDS

**Supplemental Table 1**.

*Datasets used in this study*. Statistics for the datasets generated in this study (MNase-ChIP-seq, ATAC-seq and chromatin-associated-RNA-seq) and for publicly available TF ChIP-seq profiles re-analyzed as part of this study. FRIP = Fraction of Reads in Peaks.

**Supplemental Table 2**.

Genomic coordinates of enhancers and symmetry. For each enhancer element, its symmetric /asymmetric classification is provided. The strand information encodes the position of the strongest MNase-ChIP-seq signal with respect to the enhancer core (‘+’ = downstream, ‘–’ = upstream). The mappability column highlights elements exhibiting good average mappability (> = 0.85) on both sides of the Pu.1 summit (± 1.5 kbp).

**Supplemental Table 3**.

*Remodeling events at promoters of LPS-inducible genes*. For each gene, early *vs* late and PRG *vs* SRG classification is indicated. A remodeling event (either a decrease in correlation, i.e. a shift, or a local decrease in signal, i.e. an eviction) occuring at the gene promoter is indicated with 1. Events are split by location (US = upstream; DS = downstream) and time point of stimulation.

**Supplemental Table 4**.

*Transcription factor binding and nucleosome remodeling events at LPS-inducible gene promoters*. For each TF and class of binding (either basal, LPS-induced or reduced, at different time points) the observed minus expected fraction of promoters in the indicated class showing remodeling are indicated, along with the corresponding *q*-value.

**Supplemental Table 5**.

*Remodeling events at H3K4me1+ accessible regions*. For each region, a remodeling event (a decrease in correlation or shift) is indicated with 1. Events are split by location (US = upstream; DS = downstream) and time point of stimulation.

**Supplemental Table 6**.

*Remodeling events at LPS-modulated TF-binding sites*. For each region, a remodeling event (a decrease in correlation, i.e. a shift) occurring at the indicated time point is indicated with 1. In addition, each region is annotated with respect to the binding of each TF, time point, and LPS modulation (either induced or reduced, as separately provided).

**Supplemental Table 7**.

*Nucleosome remodeling at sites of increased or reduced transcription factor binding at different time points after LPS stimulation*. For each TF and time point (TP), the size of the set, and the fraction and total number of remodeled sites is indicated. Data for either the sites showing induced or reduced binding upon LPS stimulation, separated in TSS-proximal or TSS-distal sites, are separately provided.

## SUPPLEMENTAL MATERIALS AND METHODS

### TF-ChIP-seq data analysis

For each ChIP-seq experiment, *de novo* motif discovery was run on the top 1,000, summit-centered regions using *findMotifsGenome.pl* from HOMER (v4.10.3)(Heinz et al. 2010), with the following parameters: −size −150,150 −len 8,10,12,14.

### Characterization of symmetric and asymmetric nucleosomal arrays at putative enhancers: repeat content and mappability biases

The complete RepeatMasker tracks (http://www.repeatmasker.org/) were downloaded from the UCSC genome browser. Intervals were split by class. Mappability tracks at single base-pair resolution for 50-mers were also downloaded from the UCSC genome browser for the mm9 reference genome. Spatially resolved signals were extracted as previously described (Barozzi et al. 2014), using a bin of 10 base pairs. Overall coverages were instead calculated as previously indicated in the main methods. Downstream functional enrichment analyses performed either on the full sets of asymmetric and symmetric sites or on those showing good average mappability (> = 0.85) on both sides of the Pu.1 summit (± 1.5 kbp) led to fully overlapping results.

### Characterization of symmetric and asymmetric nucleosomal arrays at putative enhancers: functional characterization

A number of features were tested for difference between the two groups (or among three groups when further splitting the asymmetric sites if displaying the same or the opposite orientation compared to the TSS of the nearest active gene):

- Distance to the nearest boundary of a topologically associated domain (TAD); asymmetric sites tend to be closer to a boundary, but the magnitude of the difference is small (median: 204.1 kbp compared to 207.6 of symmetric sites; *p*-value = 0.048, two-sided Wilcoxon rank-sum test).
- Distance to the nearest active gene (FPKM >=1); symmetric sites tend to be slightly closer to TSS of annotated genes (median: 81 kbp compared to 72.6; *p*-value = 0.0078, two-sided Wilcoxon rank-sum test). When considering the further splitting of the asymmetric sites, no significant difference was found between the two sub-groups (*p*-value = 0.63, two-sided Wilcoxon rank-sum test).
- Level of transcription of the nearest active gene (FPKM >=1); each region was assigned to the nearest active gene if found within a certain window from the gene’s TSS. Considering the classification into three groups, and a range of windows (5, 10, 20, 50, 100, 200, 400; kbp), no statistically significant difference was found (*p*-value range 0.3-0.89, Kruskal–Wallis test).
- Level of transcription of the intervening region between the putative enhancer and the nearest active gene; no difference was found between the three groups (chromatin-RNA; *p*-value = 0.88 for untreated and 0.9 for 4 hrs LPS, Kruskal–Wallis test).
- Level of transcription of the region itself (+/- 1.5 kbp from Pu.1 summit); both groups show a very low level of transcription (median <0.01 FPKM; chromatin-RNA, untreated) so this was not tested.

Evolutionary conservation; symmetric sites show a slightly higher fraction of regions lifting to the hg19 human reference genome compared to the asymmetric ones (62% *vs* 58%; *p*-value = 4.2e-4, Fisher’s Exact Test). In line with this, symmetric sites showed a slightly higher average *phastcons* score (placental mammals; +/- 150 bps from the Pu.1 summit) compared to the asymmetric ones (0.16 *vs* 0.14).

